# *POU6F2* mutation identified in humans with pubertal failure shifts isoform formation and alters GnRH transcript expression

**DOI:** 10.1101/2022.10.12.511883

**Authors:** Hyun-Ju Cho, Fatih Gurbuz, Maria Stamou, Leman Damla Kotan, Stephen Matthew Farmer, Sule Can, Miranda Faith Tompkins, Jamala Mammadova, S. Ayca Altincik, Cumali Gokce, Gonul Catli, Fuat Bugrul, Keenan Bartlett, Ihsan Turan, Ravikumar Balasubramanian, Bilgin Yuksel, Stephanie B. Seminara, Susan Wray, A. Kemal Topaloglu

**Affiliations:** Cellular and Developmental Neurobiology Section, National Institute of Neurologic Disorders and Stroke, National Institutes of Health; Bethesda, Maryland, 20892, USA; Division of Pediatric Endocrinology, Cukurova University, Faculty of Medicine. Adana, 01330, Turkey; Harvard Reproductive Sciences Center, The Reproductive Endocrine Unit and The Endocrine Unit of the Department of Medicine, Massachusetts General Hospital, Boston, MA, USA; Division of Pediatric Endocrinology, Health Sciences University, İzmir Tepecik Training and Research Hospital; İzmir, 35020, Turkey; Division of Pediatric Endocrinology, Ondokuz Mayis University, Faculty of Medicine; Samsun, 55139, Turkey; Division of Pediatric Endocrinology, Pamukkale University, Faculty of Medicine; Denizli, 20070, Turkey; Division of Endocrinology, Mustafa Kemal University, Faculty of Medicine; Hatay, Turkey; Division of Pediatric Endocrinology, Selcuk University, Faculty of Medicine, Konya, 42130, Turkey; Department of Pediatrics, Division of Pediatric Endocrinology, University of Mississippi Medical Center, Jackson, Mississippi, 39216, USA; Department of Neurobiology and Anatomical Sciences, University of Mississippi Medical Center, Jackson, Mississippi, 39216, USA

**Author notes:** Corresponding authors Susan Wray, A. Kemal Topaloglu. These authors contributed equally to this work (Hyun-Ju Cho, Fatih Gurbuz, Susan Wray, A. Kemal Topaloglu).

## Abstract

Idiopathic hypogonadotropic hypogonadism (IHH) is characterized by absent pubertal development and infertility, often due to gonadotropin-releasing hormone (GnRH) deficits. Exome sequencing of two independent cohorts of IHH patients identified 12 rare missense variants in *POU6F2. POU6F2* encodes two distinct isoforms. In mouse, pituitary and gonads expressed both isoforms, but only isoform1 was detected in GnRH cells. Although the function of isoform2 is well known, using bioinformatics and cells assays on a human-derived GnRH cell line, we demonstrate isoform1 can also act as a transcriptional regulator, decreasing *GNRH1* expression. The impact of two *POU6F2* variants (MT1 and MT2) was then examined. MT1, but not MT2, reduced transcriptional activity of either isoform, preventing Hes5 promoter activation by isoform2 and repression of GnRH transcripts by isoform1. GnRH transcription increases as the cells migrate into the brain. Augmentation earlier can disrupt normal GnRH cell migration, consistent with POU6F2 variants contributing to IHH pathogenesis.

## INTRODUCTION

Idiopathic Hypogonadotropic Hypogonadism (IHH) is a rare genetic disorder characterized by complete or partial pubertal failure caused by gonadotropin-releasing hormone (GnRH) deficiency. According to olfactory function, IHH is divided into two major forms, normal sense of smell (normosmic IHH, nIHH) and inability to smell, anosmia, defined as Kallmann syndrome (KS). Although nearly 50 genes have been reported to be associated with IHH(Howard & Dunkel, 2019; Louden et al., 2021), they account for only 50% of all cases indicating that other associated genes remain to be discovered. Delineating new genes involved in the development and/or function of GnRH neurons is relevant for understanding the basis of IHH pathogenesis in humans. We identified missense variants in *POU6F2* in IHH patients. Most POU-family members have known roles as transcriptional regulators, with many of them controlling cell type-specific differentiation pathways(Andersen & Rosenfeld, 2001; Kim, Han, Kim, & Schöler, 2021). Several POU domain-containing gene products modulate development, expression, and function of GnRH neurons(Leclerc & Boockfor, 2005; Wierman et al., 1997; Wolfe, Kim, Tobet, Stafford, & Radovick, 2002). POU6F2 has two distinct isoforms, with isoform2 being a known transcriptional regulator while the function of isoform1 is unclear. In this communication, we present evidence that POU6F2 isoform1 can also function as a transcription factor repressing *GnRH1* expression and that one of the POU6F2 variants identified alters isoform splicing and reduced transcriptional activity of either isoform. Together, these data are consistent with mutations in *POU6F2* contributing to the pathogenesis of IHH.

## RESULTS

Twelve rare missense *POU6F2 variants* (HGNC: 21694) in 15 patients from 12 unrelated families were identified. The pedigrees with clinical phenotypical features are depicted in Figure 1 and Table 1. Molecular genetic characteristics of the variants are shown in Table 2. Three POU domain variants (MT1,MT2,MT8) reside in regions necessary for proper protein function or dimerization (Figure 2A). The remaining variants (MT3-7) are in the transactivation domain. Ten of the 12 variants had CADD scores >20 and were either not seen in the largest reference population database (gnomAD) or occurred at an extremely rare minor allele frequency <0.0005. However, MT2 was found to be significantly more common in the newly published Turkish Variome at 0.002(Kars et al., 2021) (Table 2). No variant was previously reported in ClinVar. All were classified as variants of uncertain significance (VUS), except MT4 and MT7 which were categorized as ‘likely pathogenic” by ACMG/AMP classification(Richards et al., 2015). However, Polyphen-2(Adzhubei et al., 2010) and SIFT(Kumar, Henikoff, & Ng, 2009), two well-validated *in silico* prediction programs indicated most of these variants to be harmful (Table 2).

**Figure 1.**
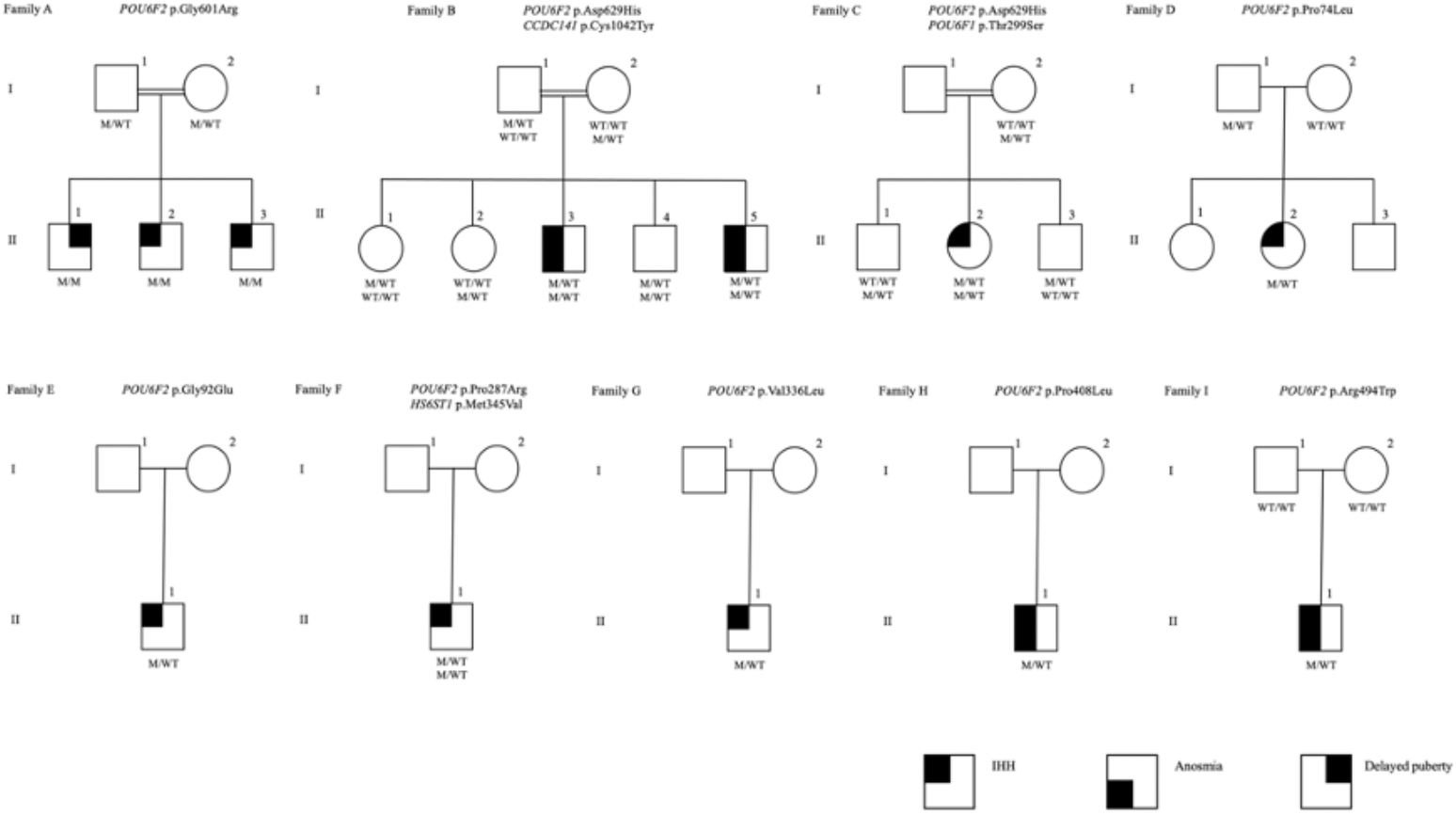
The pedigrees of the families with POU6F2 mutations. Affected males and females are represented by black squares and black circles respectively. White square symbols indicate unaffected male family members, white circle symbols represent unaffected female family members, and the double line indicates consanguinity. Under each symbol are the genotypes in the same order as the gene and variant descriptions, with WT and M denoting wild type and mutant, respectively. The legend denotes phenotypes as IHH, Anosmia, and Delayed puberty.

**Figure 2.**
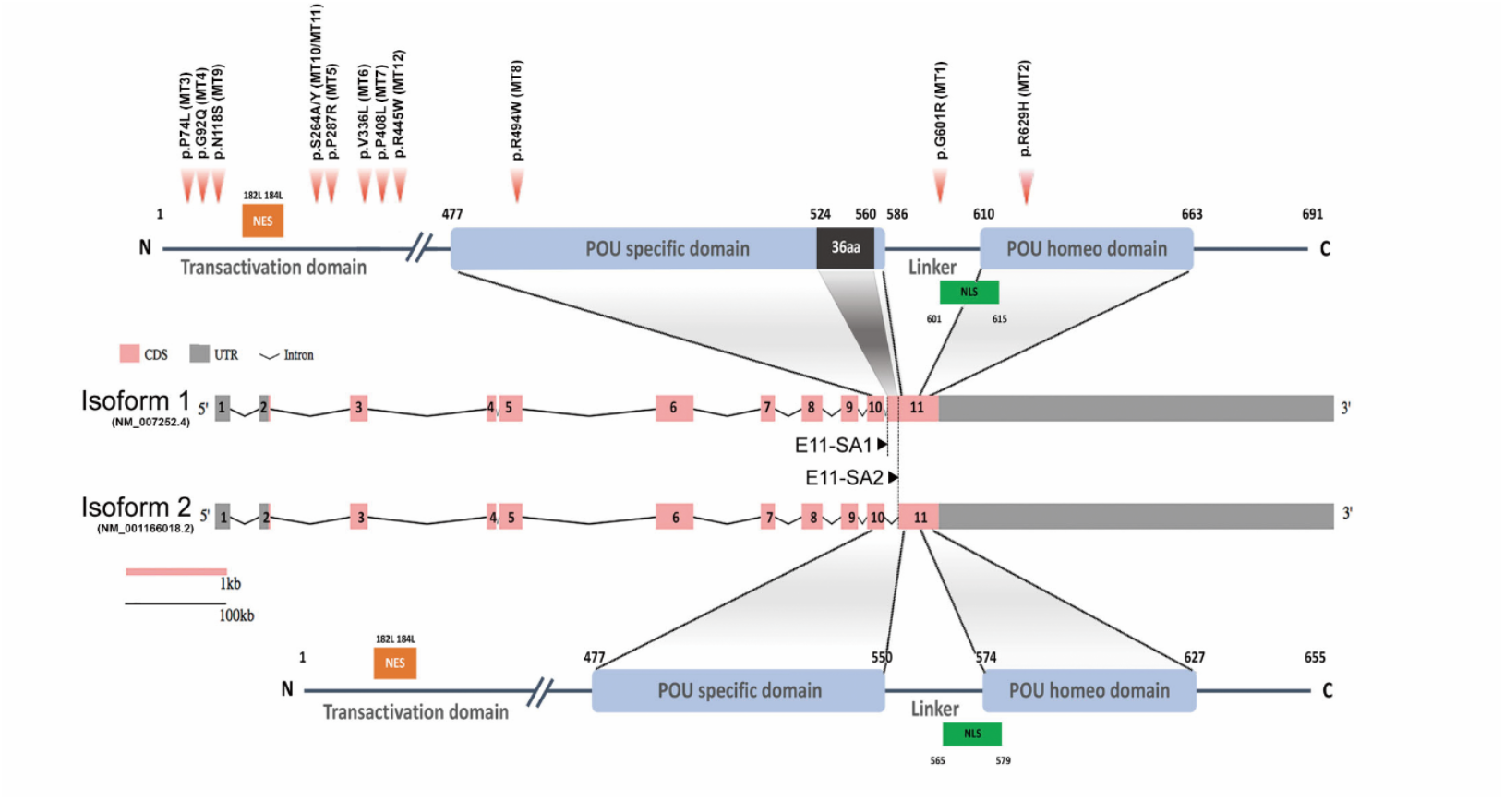
Schematic diagram of human *POU6F2* isoforms. Exon-intron structure of human *POU6F2* isoforms (middle two schematics) drawn to scale using the Gene Structure Display Server (GSDS 2.0, http://gsds.gao-lab.org). Exons are indicated by boxes to highlight the coding sequence (CDS, pink) and untranslated region (UTR, gray). Introns are indicated by black lines with a shrinked scale (0.01 ratio to scale of exons). Exon11 is alternatively spliced via two splicing acceptor sites, E11-SA1 and E11-SA2, to generate isoform1 (upper schematic) and isoform2 (lower schematic), respectively. The two conserved DNA binding domains are indicated by blue boxes and aligned to exons (encoded by exon10 to 11). Isoform1 has a unique 36aa insertion on POU specific domain (black box) not found in any other POU protein family members. The amino acid numbers are shown at the start and end point of functional domains. Twelve mutations identified from IHH patients are indicated by red arrow heads (upper schematic). Mutation 1(MT1; c.1801G>A, p.G601R in isoform1; c.1693G>A, p.G565R in isoform2) is in the linker region between the two DNA binding domains. MT 2 (c.1885A>C, p.N629H in isoform1; c.1777A>C, p.N593H in isoform2) is in the POU homeodomain. MT3-7, 9-12 are in the Transactivation domain. MT8 (c.1480C>T, p.R494W) is in the POU-specific domain. Orange boxes; Nuclear export signal (NES), green boxes; Nuclear localization signal (NLS).

**Table 1.** Clinical characteristics of individuals with POU6F2 mutations.

| Family/individual no | Mutation | Age at diagnosis (year) | Sex | Ethnicity | Initial basal LH (mIU/mL) /Estradiol (ng/dL) or Testosterone (ng/dL) | Stimulated maximum LH (mIU/mL) | Olfaction | Reproductive phenotype |
| --- | --- | --- | --- | --- | --- | --- | --- | --- |
| A II-1 | p.Gly601Arg | 15 | M | Turkish | NA | NA | Normosmic | Delayed puberty, Constitutional delay in growth and puberty |
| A II-2 | p.Gly601Arg | 18 | M | Turkish | NA | NA | Normosmic | Absent puberty, Infertility |
| A II-3 | p.Gly601Arg | 14 | M | Turkish | 0.14/<10 | 0.98 | Normosmic | Absent puberty |
| B II-3 | p.Asn629His | 15 | M | Turkish | <0.1/<10 | 3.3 | Anosmic | Absent puberty, cryptorchidism |
| B II-5 | p.Asn629His | 0.5 | M | Turkish | <0.1/<10 | 0.78 | NA | Absent mini puberty, microphallus 1 cm, cryptorchidism |
| C II-2 | p.Asn629His | 17 | F | Arabic | 0.2/0.4 | 0.2 | Normosmic | Absent puberty, primary amenorrhea |
| D II-2 | p.Pro74Leu | 16 | F | Turkish | 0.1/1.1 | 0.1 | Normosmic | Absent puberty, primary amenorrhea |
| E II-1 | p.Gly92Glu | 16 | M | Turkish | 0.1/NA | <10 | Normosmic | Absent puberty |
| F II-1 | *p.Pro287Arg* | 17 | M | Turkish | <0.1/<12 | 3.6 | Normosmic | Absent puberty |
| G II-1 | p.Val336Leu | 18 | M | American Caucasian | NA | NA | Normosmic | Absent puberty |
| H II-1 | p.Pro408Leu | 18 | M | American Caucasian | NA | NA | Anosmic | Absent puberty, cryptorchidism |
| I II-1 | p.Arg494Trp | 18 | M | Ashkenazi Jew | NA | NA | Anosmic | Absent puberty |
| J II-1 | p.Asn118Ser | 20 | M | Turkish | NA | NA | Normosmic | Absent puberty |
| K II-1 | p.Ser264Ala<br>p.Ser264Tyr | 35 | M | Ashkenazi Jew | NA | NA | Normosmic | Absent puberty |
| L II-1 | p.Arg445Trp | 18 | M | Asian | NA | NA | Anosmic | Absent puberty, microphallus at birth, cryptorchidism |

**Table 2.** The molecular genetic characteristics of the *POU6F2* variants.

| Family /individual no | Variant name | Variant at cDNA level | Variant at protein level | TRV AC | TRV MAF | GnomAD MAF | CADD score | GERP | PP2 | SIFT | ACMG/ AMP | Other IHH gene mutation /zygosity |
| --- | --- | --- | --- | --- | --- | --- | --- | --- | --- | --- | --- | --- |
| A II-1, II-2, II-3 | MT1 | c.1801G>A | p.Gly601Arg | 1 het /5170 | 0.000177 | 0.000017 | 28.8 | 5.28 | D | D | VUS: PM1, PP2, PP3 | None |
| B II-3, II-5 | MT2 | c.1885A>C | p.Asn629His | 16 hets /6724 | 0.0023 | 0.000439 | 23.6 | 2.81 | D | D | VUS: PM1, PP2, PP3 | *CCDC141* p.Cys1042Tyr Het |
| C II-2 | MT2 | c.1885A>C | p.Asn629His | 16 hets /6724 | 0.0023 | 0.000439 | 23.6 | 2.81 | D | D | VUS: PM1, PP2, PP3 | *POU6F1* p.Thr299Ser Het |
| D II-2 | MT3 | c.221C>T | p.Pro74Leu | 0 | 0 | 0.000021 | 25.1 | 5.84 | D | D | VUS: PP2 | None |
| E II-1 | MT4 | c.275G>A | p.Gly92Glu | 0 | 0 | 0.000007 | 27.8 | 5.84 | D | D | LP: PM2, PP2, PP3 | None |
| F II-1 | MT5 | c.860C>G | p.Pro287Arg | 0 | 0 | - | 25.7 | 4.24 | D | D | VUS: PM2, PP2 | *HS6ST1* p.Met345Val Het |
| G II-1 | MT6 | c.1006G>C | p.Val336Leu | 1 het /5174 | 0.000177 | 0.000027 | 23.8 | 6.17 | D | T | VUS: PM1, PP2 | None |
| H II-1 | MT7 | c.1223C>T | p.Pro408Leu | 0 | 0 | 0.000011 | 31.0 | 5.62 | D | D | LP:<br>PM1,<br>PM2,<br>PP2,<br>PP3 | None |
| I II-1 | MT8 | c.1480C>T* | p.Arg494Trp | 0 | 0 | 0.000020 | 34.0 | 5.48 | D | D | VUS:<br>PM1,<br>PP2,<br>PP3 | None |
| J II-1 | MT9 | c.353A>G | p.Asn118Ser | 0 | 0 | - | 19.9 | 5.02 | T | D | VUS:<br>PM1,<br>PM2,<br>PP2 | None |
| K II-1 | MT10 | c.790T>G | p.Ser264Ala | 0 | 0 | 0.000024 | 16.2 | 2.31 | T | D | VUS:<br>PM1,<br>PM2,<br>PP2 | None |
|  | MT11 | c.791C>A | p.Ser264Tyr | 0 | 0 | 0.000024 | 23.9 | 4.66 | D | D | VUS:<br>PM1,<br>PM2,<br>PP2 |  |
| L II-1 | MT12 | c.1333C>T | p.Arg445Trp | 0 | 0 | 0.000032 | 26.7 | 5.70 | D | D | VUS:<br>PM1,<br>PP2 | None |

Abbreviations: Het, heterozygous; AC, allele count; MAF, minor allele frequency; TRV, Turkish Variome; GnomAD, The Genome Aggregation Consortium; InterVar, Interpretation of genetic variants by the ACMG/AMP 2015; VUS, variant uncertain significance;
LP, likely pathogenic; PM, pathogenic moderate; PP, pathogenic supporting; CADD, Combined Annotation Dependent Depletion; GERP, Genomic Evolutionary Rate Profiling. Variants are described according to the RefSeq numbers following the gene names: *POU6F2*, NM\_007252; *CCDC141*, NM\_173648; *POU6F1*, NM\_001330422; *HS6ST1*, NM\_004807. PolyPhen-2, Polymorphism Phenotyping v2; SIFT, Sorting Intolerant From Tolerant; D, deleterious; T, tolerated. \* A *de novo* mutation.

In Family-A, three brothers born from a consanguineous union presented with pubertal impairment implicating an autosomal recessive mode of inheritance. All three brothers carried a homozygous mutation (p.Gly601Arg). The two younger siblings had complete IHH. The oldest sibling received monthly testosterone injections at 15yrs because of pubertal delay and by age 17 had started puberty. As different from his brothers, the milder reproductive phenotype of this patient is consistent with constitutional delay in growth and puberty, also known as self-limited delayed puberty. It has been previously observed that variants in IHH genes can also cause self-limited delayed puberty, even sometimes within the same kindreds, indicating self-limited delayed puberty shares an underlying pathophysiology with IHH(Saengkaew et al., 2021; Zhu et al., 2015). The pattern of inheritance in Family-A is clearly autosomal recessive (Fig 1). MT8 (p.Arg494Trp) in Family-I arose *de novo*. A perfect segregation of an autosomal recessively inherited mutation with pubertal impairment phenotype in multiplex families such as in Family-A was given high scores in the Clinical Genome Resource (ClinGen) framework to define and evaluate the validity of gene-disease pairs across a variety of Mendelian disorders(Strande et al., 2017). Likewise, the *de novo* variant in Family-I provides strong genetic evidence supporting causality of mutations in novel gene-disease associations(Strande et al., 2017).

The inheritance in the other pedigrees is consistent with autosomal dominant with variable penetrance and expressivity, a phenomenon commonly observed in IHH(Bouilly et al., 2018; Louden et al., 2021; Xu et al., 2017). The male patients in Family-B, Family-H, and Family-L had cryptorchidism, indicating severe congenital hypogonadism. In congenital IHH, fetal pituitary gonadotropin secretion is low, leading to inadequate fetal serum testosterone levels. As the testicular descent and growth of phallus are androgen-dependent during fetal and neonatal periods, boys with severe IHH present with micropenis, cryptorchidism, and hypospadias at birth(Pitteloud et al., 2002). The younger patient in Family-B was diagnosed with IHH based on hypogonadal features plus prepubertal gonadotropins and testosterone level at two months of age, a time window known as minipuberty, a poorly understood transient activation of the hypothalamic-pituitary-gonadal (HPG) axis between 2-6 months of age. With an appropriate physical examination and laboratory findings, it is possible to make a diagnosis of IHH during this very early window of human life(Renault et al., 2020).

In Family-C the 17yr female proband has the same mutation as the one in Family-B. In addition, she carries a distinct rare variant in another POU-family gene, *POU6F1* (See Discussion for detailed assessment). The probands in the remaining eight families (other than in families A, B, C and H) had variants in the non-POU domain part of the gene (Figure 2), the function of which remain poorly defined. We did not perform functional studies on these non-POU domain variants. However, these extremely rare variants were predicted to be deleterious by *in silico* analysis (see Table 2).

Three mutations (MT1, MT2 and MT8) are located in the POU-specific domain (POU_S_, MT8), the linker region between the POU_S_ and the POU-homeodomain (POU_H_, MT1) or in the POU_H_ (MT2, Figure 2A)(Fiorino et al., 2016; Zhou, Yoshioka, & Nathans, 1996). R494(MT8) is in the first alpha helix of the POU_S_ domain which is highly conserved in orthologs and conserved among paralogs as positively charged amino acids R or K. As such a mutation changing R to W may alter structure of this alpha helix. However, data from other POU-family members indicate that it is residues of the third alpha helix in the POU_S_ domain that are involved in hydrogen bonding with DNA base pairs(Pereira & Kim, 2009). As such we performed *in silico analysis* (Supplementary Table 1, Supplementary Figure 2), but not functional studies of MT8. In contrast to MT8, R601(MT1) and N629(MT2) are located at/or close to the edge of alpha helixes that compose the POU_H_ domain, and these are less conserved among paralogs but well conserved in orthologs. Notably, MT1 and MT2 are on exon11 which is alternatively spliced and distinguishes the two POU6F2 isoforms that have been identified (Figure 2).

### Pou6f2 isoforms are differentially expressed in mouse hypothalamic-pituitary-gonadal axis tissue

To determine which isoform might be pertinent to patients exhibiting IHH, the expression of POU6F2 isoforms in HPG axis relevant tissues was performed on mouse tissue using RT-PCR (Figure 3). *Pou6f2* is well conserved between human and mouse, except with respect to the 5’ UTR, which is located on exon1 and exon2 in human, while mouse *Pou6f2* has a shorter 5’ UTR with its coding region starting from exon1 (Compare Figure 2A, human and 3A, mouse). Thus, mouse *Pou6f2* has 9 exons which correspond to exon3-11 in the human. To date, only one *Pou6f2* mRNA sequence has been catalogued in NCBI, however, the alternative splicing of the last exon was analyzed in mouse retina cDNA and revealed the presence of both isoform1 and 2(Fiorino et al., 2016). In brain, pituitary and gonads (Figure 3B), both isoforms were present, though expression of isoform1 was more abundant than isoform2. Analysis of primary GnRH cells (Figure 3C, N=5) with primers that detect both isoform1 and isoform2, showed only isoform1 transcript present in 2 of the tested single GnRH cells. To ensure that isoform2 was not being missed due to low expression, the samples were rescreened with isoform2 only specific primers (Figure 3C). After 45 cycles of amplification, no isoform2 transcripts were detected in any of the GnRH cells, though brain cDNA was positive. Thus, in primary GnRH cells isoform1 is the predominant *Pou6f2* isoform that is expressed.

**Figure 3.**
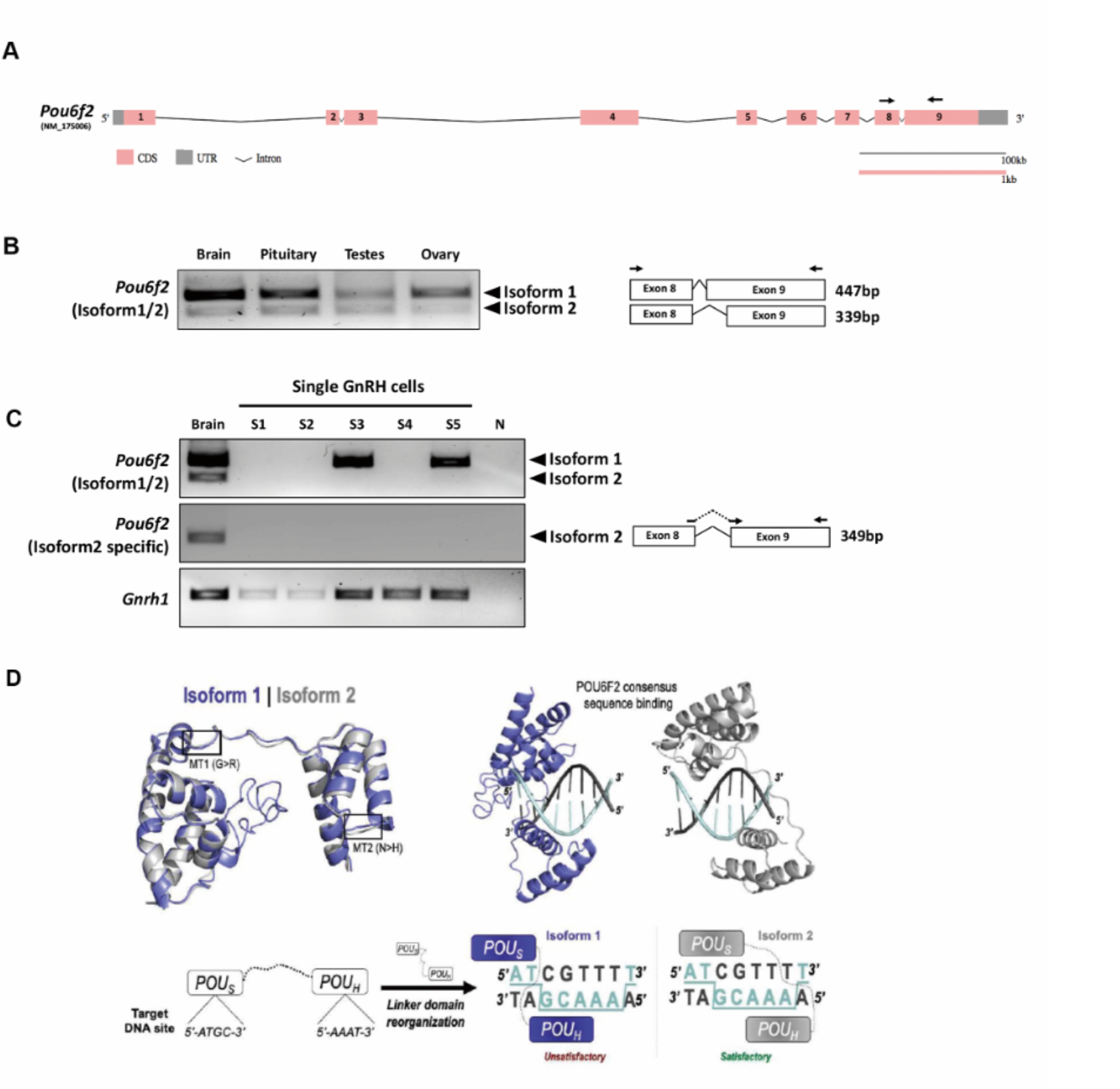
Expression of *Pou6f2* isoforms in mouse and Bioinformatic prediction of POU6F2 isoforms bound to a DNA octamer. **(A)** Exon-intron structure of mouse *Pou6f2* (GSDS 2.0, http://gsds.gao-lab.org). In mice, only one isoform has been reported which is composed of 9 exons and corresponds to isoform1 of human *POU6F2*. Primers used for PCR are shown as arrows on exon8 and 9. **(B)** Gel image of RT-PCR analysis performed in mouse tissue. Top band (447 bp) shows isoform1 and bottom band (339bp) shows isoform2 which is skipping 108bp by alternative splicing on exon9. **(C)** Gel image of RT-PCR analysis of *Pou6f2* isoforms (top and middle gel) in 5 GnRH single cells (bottom gel). Only isoform1 was detected. **(D)** Upper Left, Superimposition of isoform1 (purple) and 2 (gray) structures predicted by C-I-TASSER. The location of MT1 and MT2 is indicated by boxes. Upper Right, HDOCK prediction of POU6F2 binding to the OCT1 DNA consensus site (5’-ATGCAAAT-3’). Template-free docking was used to prevent simulation bias. Lower Left and Right, Structural representation of the interaction between each isoform and dsDNA octamers. Two-dimensional cartoon illustrating the molecular interactions between each POU domain and their predicted binding sites. Satisfactory (for isoform2) and unsatisfactory (for isoform1) binding modes are indicated. **Figure 3-source data B and C**. Raw images of uncropped and cropped gels from panels B and C are included in the zipped source data as original tiffs and annotated pdf format.

### POU6F2 modeling

Previous studies have shown that POU6F2 isoform2 binds to divergent POU and combined SOX/POU DNA sequences and mediates transcriptional activity while isoform1 binds poorly to these sequences and shows little transcriptional activity(Fiorino et al., 2016; Zhou et al., 1996). *In silico* analysis was used on the two isoforms to establish the validity of our modeling. Although POU6F2 is yet to be crystallized, a closely related paralog human POU6F1 has been crystallized bound to an octamer motif(Pereira & Kim, 2009) which increases the accuracy of homology modeling(Haddad, Adam, & Heger, 2020). C-I-TASSER produced structures for POU6F2 isoform1 and 2 (Figure 2B) using the POU6F1 crystal template with good resolution (PDB code: 3D1N; resolution=2.51 Å). The POU domains for each POU6F2 isoform were in the same fold as POU6F1 (TM-score_iso1_=0.66, RMSD_iso1_=1.03; TM-score_iso2_=0.80, RMSD_iso2_=0.81). Highly variant N-terminal domains upstream of the POU domains were not detected in the original POU6F1 crystal and were disordered in POU6F2 structures and thus were omitted in downstream structural experiments. Previous experiments indicated that only isoform2 binds to OCT1 consensus DNA(Zhou et al., 1996). w3DNA was used to predict the structure of the human OCT1 DNA consensus sequence (5’-A_1_T_2_G_3_C_4_A_5_A_6_A_7_T_8_-3’), and HDOCK predicted a more favorable scoring function of binding for isoform2 (-303.26au) compared to isoform1 (-258.88au). The predicted binding mode showed that isoform2 POU_S_ binds to 5’-A_1_T_2_G_3_C_4_-3’ and POU_H_ to 5’-A_5_A_6_A_7_T_8_-3’ by embracing both faces of dsDNA, whereas isoform1 did not (Figure 3D). This is consistent with literature showing that isoform2 can bind these octamer consensus sequences, whereas isoform1 cannot(Zhou et al., 1996).

### POU domain mutation identified in IHH patients alters splicing preference of human exon11

Alternative splicing of *POU6F2* has been identified in human retina cDNA(Zhou et al., 1996), but the proportion of the mRNA isoforms derived from those splicing events is not well known. The most studied splicing event is on exon11, which distinguishes isoform1 and isoform2. Two mutations (MT1 and MT2) identified in this study reside in exon11 which can affect both isoforms (Figure 2A). To predict the possible effect of both mutations on splicing of exon11 and isoform expression, *in silico* splicing analysis was performed using online prediction programs. Both mutations are predicted to affect splicing by 3 out of 4 prediction programs (Figure 4A).

**Figure 4.**
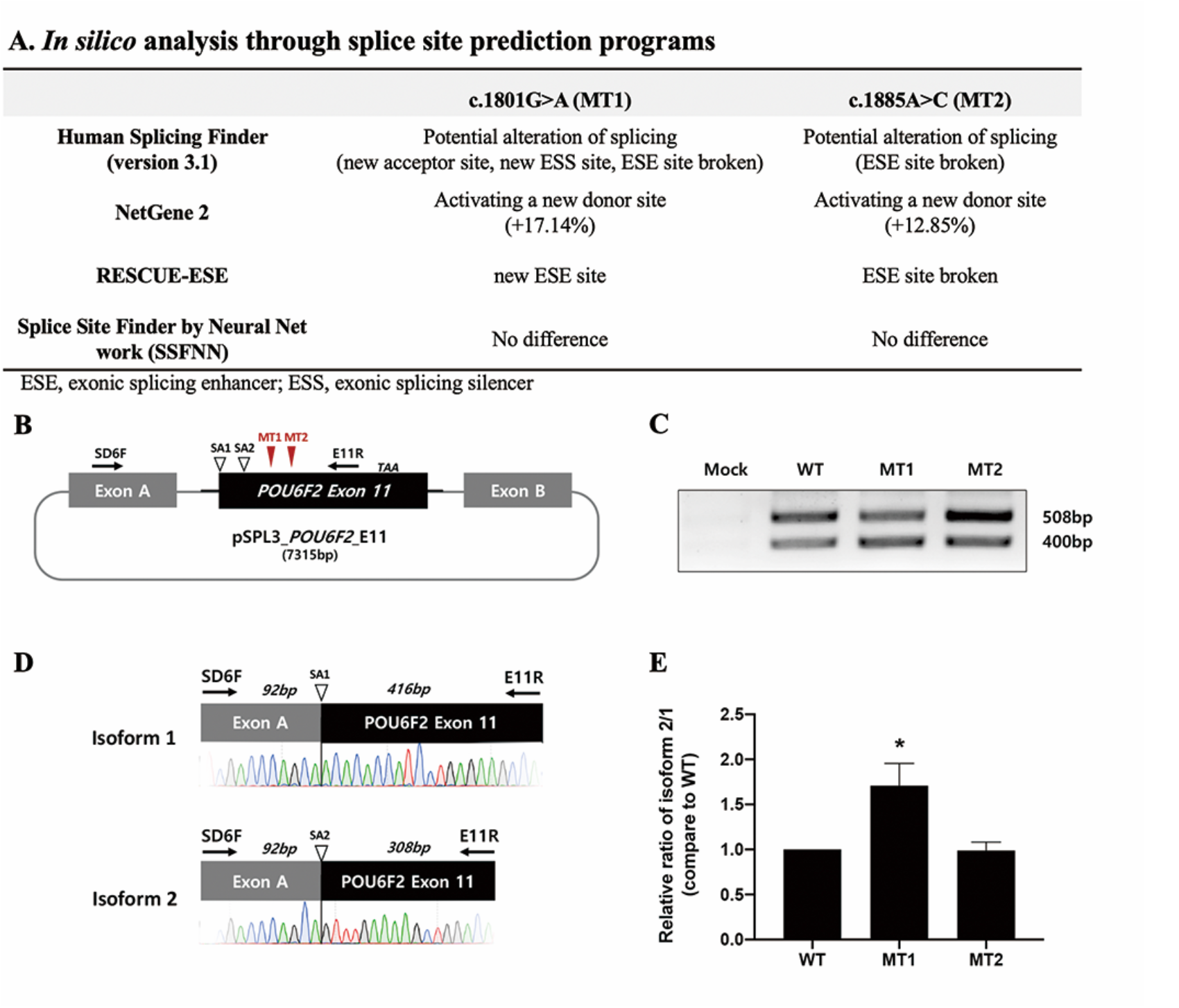
*In silico* analysis of splice sites and the effect of MT1 or MT2 on isoform expression using an *in vitro* splicing assay of human exon11. **(A)** Table showing results of MT1 and MT2 on splice sites using *in silico* analysis. Three of the four programs predicted mutational changes. **(B-E)** *In vitro* splicing assay of human POU6F2 exon11. **(B)** Exon11 of human *POU6F2* (Black box) and flanking intronic sequences (~300bp, Black line) were inserted into pSPL3 minigene vector which has two vectoral exons (ExonA and B, gray boxes). Splicing acceptor site for isoform1 (SA1) and for isoform2 (SA2) are indicated by empty arrow heads. Two identified mutations (red arrow heads) are located on the exon11 after SA2 splicing acceptor site and before the stop codon (TAA). **(C)** RT-PCR using SD6F (forward primer on exonA) and E11R (Reverse primer on POU6F2 exon11) resulted in two different sized bands (isoform1 and isoform2) in assays using WT, MT1 and MT2. **(D)** Schematic diagram shows the amplified isoform1 and isoform2 fragments from WT. Sequencing analysis confirmed the splicing junction from exonA to SA1 (isoform1) and SA2 (isoform2). **(E)** Quantitative analysis of isoform1 and 2, from WT, MT1 and MT2 constructs, performed via qPCR and represented as a relative isoform2/isoform1 ratio. MT1 increased isoform2 compared to WT (*N*=3). \**P*<0.05. **Figure 4-source data C**. The zipped source data file contains raw images of the uncropped and cropped gels from panel C (as original tiffs and annotated pdf format).

To directly analyze the possible splicing changes on exon11, *in vitro* splicing assays were performed by introducing human exon11 into pSPL3 minigene plasmids in between two artificial exons designed to be spliced into a single mRNA (Figure 4B). Plasmids were transfected into HEK293 cells, cultured for 2 days and processed mRNAs obtained. RT-PCR using SD6F (forward primer on exonA) and E11R (reverse primer on POU6F2 exon11) showed two different sized bands from both WT and mutant sequences but not pSPL3 mock (Figure 4C). The upper band (508 bp) is spliced from exonA (92bp) to exon11 (416bp) via the first splicing acceptor site (SA1) forms isoform1. The smaller band (400bp) is spliced from exonA to exon11 (308bp) via the second splicing acceptor site (SA2) forms isoform2. The DNA sequence of each band was confirmed by TA-cloning of PCR products and plasmid sequencing (Figure 4D). Neither new splicing variants nor absence of normal splicing variants were observed from either mutant sequence. However, the band intensities of the isoforms changed with the introduction of MT1. In WT, the isoform1 splicing band was stronger than the isoform2 splicing band (Figure 4C). Introduction of MT1 reversed this relationship, with the isoform2 band now stronger, though isoform1 was still present (Figure 4C). To obtain a better measurement of the isoforms, qPCR analysis was performed using primers specific for each isoform. The ratio of isoform2/1 was calculated and represented as a relative value compared to WT (Figure 4E). MT1 resulted in a significant increase in the ratio of isoform2/1 (~1.6-fold higher than WT) while the ratio obtained for mutation 2 (MT2) was similar to WT. No significant differences were observed with or without cycloheximide treatment suggesting that these spliced mRNAs were stable from nonsense-mediated decay. To further evaluate the potential effects of MT1 and MT2 on isoform1 and isoform2, functional evaluation of the mutant proteins was performed using computational modeling and *in vitro* transcriptional assays.

### Functional analysis of POU6F2 isoform2 as a known transcription factor

POU6F2 isoform2 is a known transcriptional regulator(Fiorino et al., 2016). To examine the alterations induced by MT1 and MT2 on isoform2 (Figure 5A), DynaMut and CABS-flex were used to determine changes in protein structure. DynaMut predicted that both mutations stabilized isoform2 folding (ΔΔG_MT1_=0.956; ΔΔG_MT2_=0.211, Figure 5B). Protein flexibility dictates not only a protein’s function but also its ability to respond to stochastic environmental changes(Teilum, Olsen, & Kragelund, 2011). CABS-flex revealed that isoform2 protein flexibility was largely diminished by both mutations (P=0.0001, figure 5B). SAMPDI was then used to predict changes in the affinity of isoform2, in the setting of either MT1 and MT2, for the known OCT1 DNA consensus. Both mutations were predicted to destabilize this interaction. In sum, *in silico* analysis predicted that the MT1 and MT2 found in IHH patients would be deleterious to the intrinsic properties of the POU6F2 isoform2, including both its protein flexibility and affinity for DNA.

**Figure 5.**
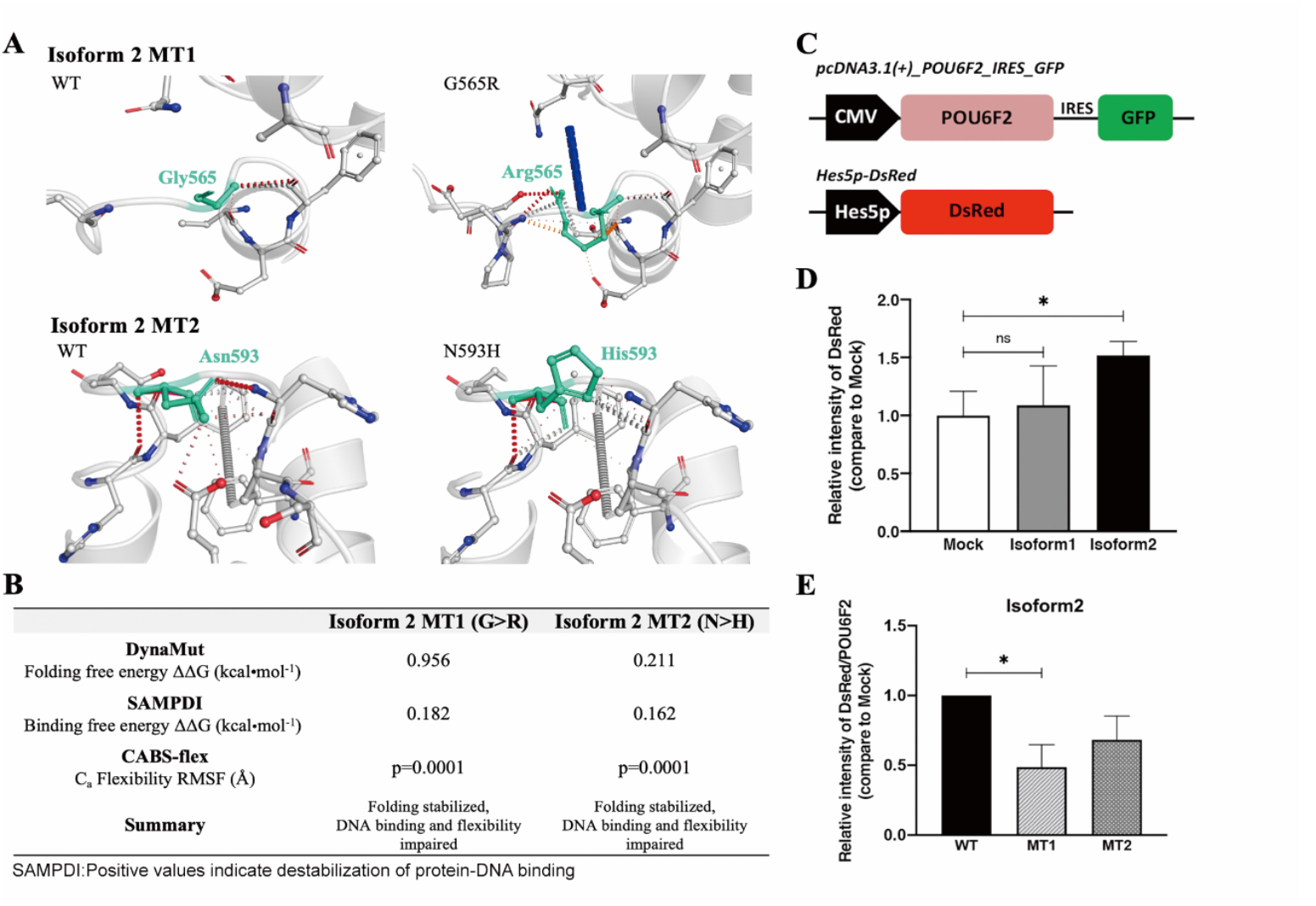
Structural analysis of IHH mutations in POU6F2 isoform2 and *in vitro* transcription assay of human POU6F2 on *Hes5* promoter. **(A)** DynaMut prediction of WT and mutant proteins for isoform2. Individual amino acid substitutions are indicated in cyan. **(B)** Structural evaluation scores indicating how MT1 and MT2 affect POU6F2 isoform2 protein folding (DynaMut), natural protein flexibility (CABS-flex) and DNA binding (SAMPDI). DynaMut and CABS-flex represent changes in the individual protein structures, whereas SAMPDI represents changes in the affinity of POU6F2 to bind the OCT1 DNA consensus site (‘5-ATGCAAAT-3’). Characterization of stabilizing or destabilizing effects are indicated. CABS-flex values analyzed using a paired-t test. **(C-E)** Transcriptional activity of POU6F2 isoforms were evaluated by *in vitro* transcription assay. **(C)** Schematic of vectors co-transfected into HEK293FT cells. For POU6F2 expression vector, the CDS of each isoform was inserted into pcDNA3.1(+)_IRES-GFP after the CMV promoter. The reporter vector included the promoter sequences (~76bp) of *Hes5* followed by DsRed (Hes5p-DsRed**). (D)** After Western blot analysis using an anti-DsRed antibody, band intensities were measured (image J software) and represented as a relative value to that of the mock vector group (bar graph). Only isoform2 showed a significant increase in DsRed expression (~1.5 fold of mock). **(E)** Using the same assay, analysis for isoform2 mutations were performed. The band intensity of DsRed is normalized by POU6F2 intensity and compared to WT. The relative values are represented in the bar graph, showing that MT1 significantly decreased transcriptional activity (~0.5 fold of wild type). ns, not significant; \**P* <0.05 **Figure 5-source data D and E**. The zipped source data file contains: 1) statistical data represented in panel D with and without normalization to WT control groups (pdf format), and 2) raw data and blots used for quantification from the Hes5 reporter assay (excel format).

To directly evaluate the transcriptional activity of WT and mutant isoform2 POU6F2 proteins, *in vitro* transcription assays were performed using a DsRed reporter under the *Hes5* promoter (760bp, Figure 5C). *Hes5* is known to be upregulated by isoform2 overexpressed in HEK293 cells(Fiorino et al., 2016) and the promoter sequence used for assay includes two predicted POU protein binding sequences (5’-AAGCAAAT-3’ and 5’-ATGCTAAT-3’; predicted by PROMO v3.0.2). Isoform1 (non-*Hes5* DNA binding control) or isoform2 (DNA-binding) expression vectors were co-transfected into HEK293FT cells with Hes5p-DsRed plasmids; then the expression of DsRed was analyzed by Western blot assay. Isoform2 showed a significant increase in DsRed expression (~1.5-fold over mock) while isoform1 was similar to mock (Figure 5D). These data are consistent with only isoform2 acting as a transcription factor protein for *Hes5* POU protein binding sequences. This assay was then repeated using the mutated isoform2 variants. DsRed expression was normalized by POU6F2 and presented as a relative value compared to the WT (Figure 5E). Transcriptional activity was reduced by 50% in the MT1 group (P=0.0334) compared to WT. MT2 also showed decreased activity, but it was not statistically different from the WT group (P=0.1367).

To complement the transcription factor binding assays, HDOCK was used to predict the interaction of isoform2 with the two Hes5 POU binding sequences described above, 1) 5-<u>CCA</u>A_1_A_2_G_3_C_4_A_5_A_6_A_7_T_8_-3’, which also contains a three base pair 5’ flank upstream (underlined above) and a T>A substitution in the second position (A_2_) relative to the OCT1 consensus; and 2) 5’-A_1_T_2_G_3_C_4_T_5_A_6_A_7_T_8_-3’ which has an A>T substitution in the fifth position (T_5_) relative to the OCT1 consensus. HDOCK predicted a more favorable scoring function between isoform2 and the Hes5 POU binding site 1(-340.48au) compared to site 2(-234.19au, respectively). For site 1, POUS docked onto sense 5’-A_1_A_2_G_3_C_4_-3’ and POU_H_ onto antisense 3’-T_5_T_6_T_7_A_8_-5’; these interactions are consistent with the literature(Zhou et al., 1996). Unlike the data obtained from the *in vitro* transcription assay described above, SAMPDI (Supplementary Table 1) predicted that both MT1 and MT2 would impair DNA binding to site 1(ΔΔG_MT1_=0.271; ΔΔG_MT2_=0.548 kcal mol^-1^) or site 2 (ΔΔG_MT1_=0.261; ΔΔG_MT2_=0.254 kcal mol^-1^), i.e. decrease the POU6F2-Hes5 binding affinity for isoform2.

### Functional analysis of POU6F2 isoform1 as a potential transcription factor

Compared to POU6F2 isoform2, the function of POU6F2 isoform1 is unclear. However, using the Yeast One-Hybrid System with a 5’-upstream region of the porcine *Fshβ* as the bait sequence, Yoshida et al. (Yoshida et al., 2014), cloned a cDNA encoding a partial sequence of the POU domain from porcine pituitary. The clone was equivalent to POU6F2 isoform1 and was able to modulate expression of developmental pituitary genes, using transient transfection assays of promoter activity in CHO cells. We tested isoform1 against the predicted *Fshβ* protected site(5’-ATAAGCTTAAT-3’). Modeling showed that not only does isoform1 bind in the correct orientation (i.e., insert facing away from DNA) but the POU_S_ binds onto ATAA and POU_H_ onto TTAA, which agrees with the sites that POU6F1 monomer2 uses to bind CRH (crystal PDB code: 3D1N). Examining the *GnRH1* promoter we found a similar site but with 3 mismatches (<u>AAAA</u>GC<u>ATAG</u>T, region of *GnRH1* promoter sequence that aligned with *Fshβ*). When we tested this site using HDOCK, isoform1 did not interact with the correct domains/orientation and the docking solutions were not in agreement with the binding mode predicted with *Fshβ*. However, we noticed that the *GnRH1* promoter region contains a reverse-complementary version of the POU6F2 consensus site with one A/T substitution (POU6F2 consensus: 5’-ATGCAAAT-3’; *GnRH1* site: 5’-TACGAAAA-3’ = 3-ATGCTTTT-5’, Figure 6A). Using 3D modeling, one sees the POU6F2 consensus site arrangement and appropriate binding for isoform1 (Figure 6B, i.e. POUs to ATGC half and POUh to AAAA half), which is in fact, in agreement with isoform2. The 44-nucleotide insert on isoform1 sticks away from the complex allowing it to bind. Next, DynaMut and CABS-flex were used to determine changes in protein structure that might be induced by MT1 and MT2 on isoform1 (Figure 6C and D). DynaMut predicted that MT1 destabilized while MT2 stabilized isoform1 folding (ΔΔ_GMT1_=-0.562; ΔΔG_MT2_=1.728). CABS-flex revealed that isoform1 protein flexibility was decreased only with MT2 (P=0.0016).

**Figure 6.**
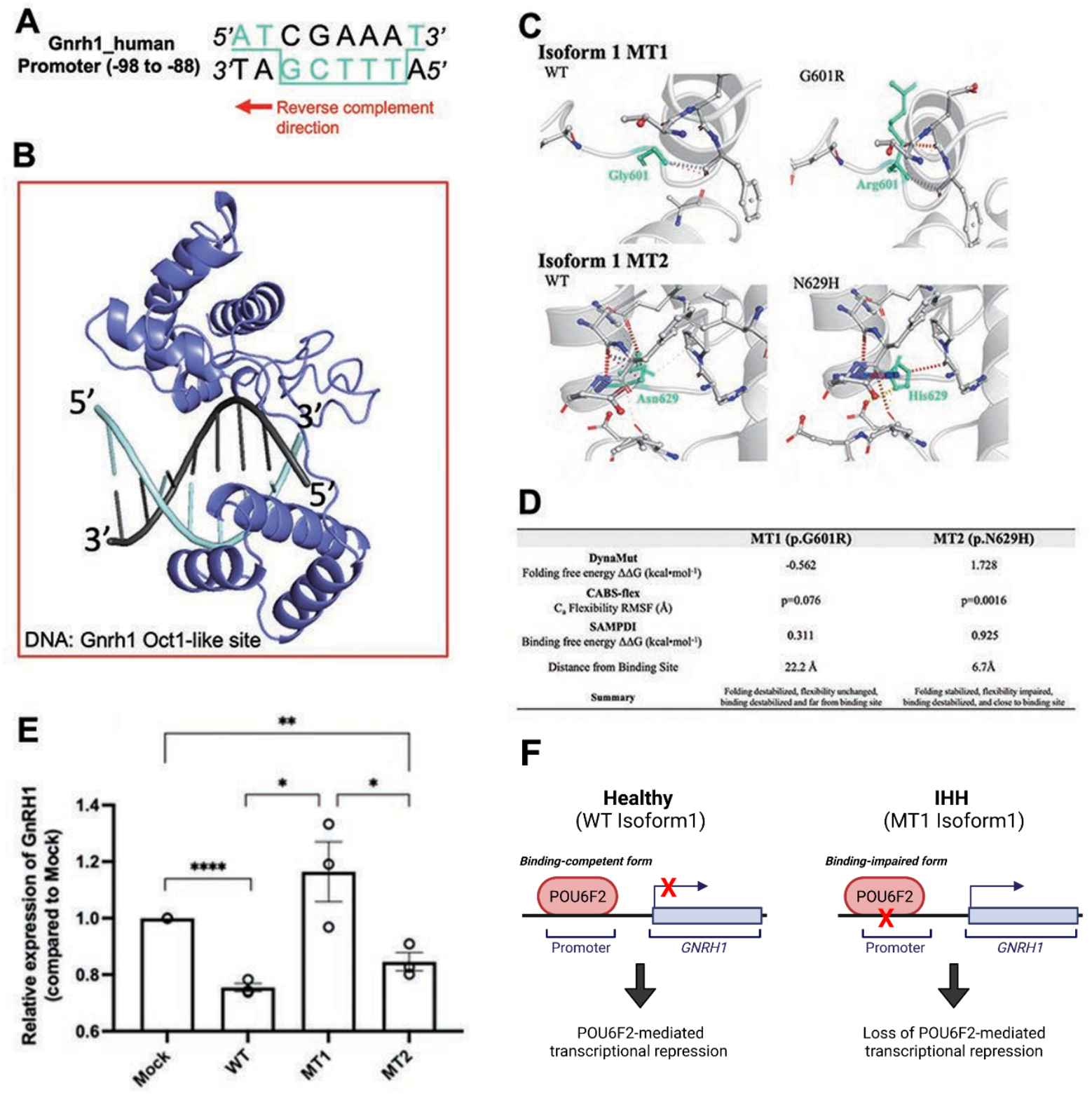
Structural analysis of IHH mutations MT1 and MT2 on POU6F2 isoform1 and *in vitro* transcription assay of isoform1 in immortalized human GnRH cells. **(A)** OCT1 consensus-like site (5’-ATGCTTTT-3’) is identified in human *GnRH1* promoter (-98 to -88). Binding site in 3D modeling uses POU_S_ to ATGC, and the POU_H_ is predicted to insert into a groove between both faces of the dsDNA, thus making contact with both TTTT and AAAA. **(B)** HDOCK prediction of POU6F2 isoform1 binding to the OCT1 consensus-like site. Template-free docking was used to prevent simulation bias. **(C)** DynaMut prediction of WT and mutant proteins for isoform1. Individual amino acid substitutions are indicated in cyan. **(D)** Structural evaluation scores indicating how MT1 and MT2 affect POU6F2 isoform1 protein folding (DynaMut), natural protein flexibility (CABS-flex) and DNA binding (SAMPDI). DynaMut and CABS-flex represent changes in the individual protein structures, whereas SAMPDI represents changes in the affinity of POU6F2 isoform1 to bind the OCT1 consensus-like site (5’-ATGCTTTT-3’). Characterization of stabilizing or destabilizing effects are indicated. CABS-flex values analyzed using a paired-t test. **(E)** Quantitative RT-PCR of *GnRH1* in FNC-B4-hTERT cells transfected by POU6F2 isoform1. The expression of *GnRH1* was normalized to each experimental mock group and the relative values are represented in the bar graph. MT1 significantly increased *GnRH1* transcript compared to both WT and MT2 groups but was not significantly different from the Mock group. **(F)** Schematic summary of isoform1 as a transcriptional regulator generated by Biorender (https://biorender.com/). Un-paired t test was performed (*N*=3), \**P*<0.05, \*\**P*<0.001, \*\*\*\**P*<0.0001 **Figure 6-source data E**. The zipped source data file contains raw data from the RT-qPCR measurements presented in panel E (excel format).

To directly evaluate the transcriptional activity of WT and mutant isoform1 POU6F2 proteins, *in vitro* transcription assays were performed using a human GnRH cell line FNC-B4-hTERT(Hu et al., 2009). GnRH transcript was measured using qPCR. Isoform1 but not isoform2 was found to be expressed in FNC-B4-hTERT cells, whereas both isoforms were detected in human brain lysate, albeit isoform2 levels were drastically lower compared to isoform1 (Supplementary Figure 1). Further, expression of either WT-POU6F2 or MT2-POU6F2 isoform1 in these cells significantly decrease GnRH expression compared to mock (Mock=1; WT=0.7547±.014, \*\*\*\**P*<0.0001; MT2=0.8458±0.032, \*\**P*<0.001; Figure 6E). No significant difference was found between WT and MT2 isoform1 treated groups (Figure 6E). Notably, MT1 significantly increased *GnRH1* transcript compared to both WT and MT2 groups (MT1=1.164±0.11, P<0.05 for both comparisons) but was not significantly different from the Mock group (Figure 6E). These experiments reveal that isoform1 POU6F2 proteins, transcriptionally regulate *GnRH1* (Figure 6F). Table 3. summarizes the *in vitro* experiments performed in this study.

**Table 3.** Summary of *in vitro* experiments.

|  | Expression in<br>primary mouse<br>GnRH neurons<br>(Fig3) | Exon11 Splicing assay in minigene<br>Transfected HEK293 cells (Fig4) |  |  | in vitro transcription assays using a DsRed<br>reporter under the Hes5 promoter in HEK293<br>cells (Fig5) |  |  | Isoform1 *in vitro* transcription assays in<br>human GnRH cell line FNC-B4-hTERT (Fig6) |  |  |
| --- | --- | --- | --- | --- | --- | --- | --- | --- | --- | --- |
|  |  | WT | MT1 | MT2 | WT | MT1 | MT2 | WT | MT1 | MT2 |
| **POU6F2**<br>**Isoform1** | Present | Normal | Decreased or<br>normal | Normal | Normal | Not Assayed | Not Assayed | Decreased | Increased | Decreased |
| **POU6F2**<br>**Isoform2** | Absent | Normal | Increased | Normal | Increased | Decreased | Decreased but<br>ns | Not Assayed | Not Assayed | Not Assayed |

## DISCUSSION

Our findings demonstrate POU6F2 is necessary for human puberty and reproduction; they also reveal a novel action of POU6F2 isoform1 as a transcription factor capable of acting on the GnRH promotor. We present clinical and molecular genetics data from 15 patients who belonged to 12 independent families that highlighted POU6F2 variants. These patients all presented with pubertal failure and were diagnosed with IHH. Two of the POU6F2 mutations which would potentially alter DNA binding were then analyzed by molecular, cellular and bioinformatic techniques. These studies indicate that one of the POU domain mutations (MT1) alters the DNA binding capacity of both POU6F2 isoform1 and isoform2, and by extension, affects transcription efficiency. The lack of an effect by MT2 is consistent with new Turkish variome data(Kars et al., 2021) (unlike that in GnomAD), indicating that this mutation is probably too common in the Turkish population to cause IHH by itself, but may contribute to the phenotype in combination with other mutations.

Since IHH patients have low gonadotropins in the face of prepubertal serum sex steroid levels, the pathophysiology of this condition should reside in the pituitary and/or hypothalamus. POU6F2 has been shown to be highly expressed in the early embryonic pituitary and stimulate the expression of PROP1(Yoshida et al., 2014), though the specific isoforms were not examined. PROP1 is well known to induce POU1F1 and the development of gonadotropes and corticotropes in the anterior pituitary(Kioussi, Carriere, & Rosenfeld, 1999). POU1F1 also induces differentiation of GH, PRL, TSHB producing cell lineages in the anterior pituitary(Andersen & Rosenfeld, 2001) and when mutated, causes multiple pituitary hormone deficiency syndrome and hypopituitarism(Turton et al., 2005). However, it is unlikely that IHH in our patients is due to impaired pituitary effects of POU6F2 via PROP1 since our patients have only a deficiency of LH and FSH and not ACTH or any of the remaining three pituitary hormones (Growth hormone, Prolactin, and TSH) induced by POU1F1 subsequent to PROP1 stimulation.

There are many modulators of reproduction within the hypothalamus, but most are translated to the pituitary-gonadal axis via GnRH neurons and dysregulation of GnRH neurons prenatally or postnatally can result in an altered HPG axis. In this report we show that WT-POU6F2-isoform1 can directly inhibit *GnRH1* transcription and that MT1 alters the transcriptional activity of both isoform1 and 2. POU6F2 isoform2 in the POU domain region including the linker sequence has a high similarity with POU6F1. The recent crystallization of POU6F1 revealed that members of the POU6-family can bind target DNA as dimers such that the POU_S_ and POU_H_ domains of one monomer bind opposite faces of dsDNA via a flexible linker region, and that the POU_S_ domain of the other monomer binds adjacent to the first POU_S_(Pereira & Kim, 2009). This is in contrast with PIT-1 (also known as POU1F1) which binds the same face of dsDNA as a dimer but shares features of OCT1 (also known as POU2F1) which binds opposite faces of dsDNA as either a monomer or dimer(Herr & Cleary, 1995; Jacobson, Li, Leon-del-Rio, Rosenfeld, & Aggarwal, 1997; Reményi et al., 2001; Ryan & Rosenfeld, 1997). Since POU6F2 is the closest related clade member to POU6F1 and the fact that members of the same subclass of POU domain factors tend to have similar DNA-binding preferences(Andersen & Rosenfeld, 2001), it is expected that they bind DNA in a comparable manner. In fact, computational modeling found that isoform2 docked onto CRH in a manner identical to monomeric POU6F1, whereas isoform1 did not (Supplementary Table 1). To validate the HDOCK template-free binding benchmark, a simulation was run to test if the server could predict POU6F1 binding to CRH promoter from the individual structures and found that the prediction was in the exact same orientation as for the POU6F1-CRH co-crystal. The spatial arrangement in combination with overlapping interfacial contacts confirms that POU6F2 isoform2 is comparable to POU6F1 DNA binding to the CRH motif.

Computational modeling also found that isoform1 did not dock onto CRH in a manner identical to monomeric POU6F1. POU6F2 isoform1 was shown to interact with a region of the *FSHβ* promoter(Yoshida et al., 2014). Although a similar region (3 nucleotide changes) was found in the GnRH promoter, modeling did not show binding. However, an OCT1 consensus-like site(5’-ATGCTTTT-3’) was identified in the human *GnRH1* promoter(-99 to -92). 3D modeling predicted that the POU_S_ bound to ATGC, and the POU_H_ inserted into a groove between both faces of the dsDNA, contacting both the TTTT and AAAA. Quantitative RT-PCR of *GnRH1* in a human cell line transfected with POU6F2 isoform1 showed exogenous expression of POU6F2 isoform1 decreased GnRH transcript levels confirming our 3D modeling.

Prenatally, GnRH cells migrate from the olfactory placode into the developing forebrain. Alterations in GNRH expression occur during migration with the cells pausing at the nasal forebrain junction(Duittoz et al., 2021). As they enter the forebrain, there is a significant increase in GnRH transcription(Simonian & Herbison, 2001) with concomitant changes in protein expression(Kramer, Krishnamurthy, Mitchell, & Wray, 2000; Kramer & Wray, 2000) as well as neuronal activity(Duittoz et al., 2021). Previous studies in mouse showed that MSX and DLX, non-Hox homeodomain transcription factors, compete for same binding site and alter GnRH transcription differently, with DLX enhancing and MSX repressing GnRH expression(Givens et al., 2005). The authors reported that MSX mutant mice had more GnRH expressing cells at E13.5, yet the majority of these cells were confined to nasal regions being distributed in both expected regions as well as ectopically in the olfactory epithelium. In addition, the study reported that the mouse GN11 cell line, a model for immature migrating GnRH cells, expressed MSX while the GT1-7 cells, a model for mature mouse GnRH cells, expressed both DLX and MSX. The human cell line used for isoform1 assays in the present study is derived from olfactory mucosa, representing an immature GnRH cell. WT-POU6F2-isoform1 may thus have a role in maintaining low GnRH transcription levels specifically when GnRH cells are outside the forebrain, prioritizing migration over gene expression. This study suggests that mutations (such as MT1) releasing POU6F2 isoform1 repression would be detrimental to the developing GnRH neuronal system resulting in IHH.

In the adult, POU6F2 is expressed in the dorsal hypothalamus in a scattered fashion(Yoshida et al., 2014; Zhou et al., 1996), which may overlap with the dispersed location of GnRH neurons in the hypothalamus(Herbison, Porteous, Pape, Mora, & Hurst, 2008). Other POU domain genes(POU3F1 also known as OCT6(Wierman et al., 1997; Wolfe et al., 2002) and POU2F1 also known as OCT1(Leclerc & Boockfor, 2005)) have been shown to repress(Wierman et al., 1997) or enhance(Leclerc & Boockfor, 2005; Wolfe et al., 2002) *GNRH1* expression. Wierman et al(Wierman et al., 1997) speculated that POU3F1 is able to turn off and on the transcriptional machinery in postnatal GnRH cells, influenced by the hormonal environment (such as sex steroids), when groups of GnRH cells were reported to be unable to express the mature gene product(King & Rubin, 1995). As an alternative/additional mechanism of disease via mechanisms post-GnRH cell migration into the forebrain, the effects of POU6F2 variants to impair pubertal development may occur indirectly to GnRH cells via the arcuate(infundibular) nucleus. The arcuate kisspeptin neurons have been proposed as the hypothalamic GnRH pulse generator driving fertility(Clarkson et al., 2017). Campbell et al. profiled gene expression in the arcuate nucleus of the hypothalamus in adult mice and found *Pou6f2* is highly expressed with a subgroup of *Pomc* neurons, a major anorectic gene, which may also give rise to kisspeptin neurons(Campbell et al., 2017; Sanz et al., 2015). In addition, a single-cell transcriptome analysis of the hypothalamic arcuate nucleus in E15 mouse showed that *Pou6f2* was one of the transcription factors showing differential expression among subclusters(Huisman et al., 2019).

It is well recognized that allele specific expression(Kukurba et al., 2014) and/or alternative splicing across tissues(Gutierrez-Arcelus et al., 2015) are important variables in determining the disease-causing potential of a missense variant. Therefore, the impact of a variant is better estimated by taking into consideration not only genome but also transcriptome data at the tissue level(Li et al., 2017). Our study showed that a point mutation in a coding region can result in missense mutant proteins across two isoforms due to splicing events. Considering the differential ratio of *Pou6f2* isoforms in mouse tissues and primary GnRH cells themselves, investigating the role of each isoform and the effect of mutations on their expression in relevant tissues is necessary to unveil the pathogenic mechanism of POU6F2 in human IHH. This is particularly essential as the central components of the HPG axis are among the least accessible tissues in human body. Our *in silico* and *in vitro* assays of POU domain mutations, show that MT1 increased the generation of isoform2 to isoform1 by altering splicing preference. This could have a profound effect on the GnRH cell subpopulation that predominantly expresses isoform1. If GnRH cells were now producing isoform2, new transcriptional targets could be impacted, disrupting the normal activity of downstream genes. Our experiments do not pinpoint whether it is this change in isoform1/isoform2 production and/or altered isoform1 activity that perturbs GnRH function *in vivo*. However, the defective function of isoform1 or a decreased amount of isoform1, could lead to increased GnRH1 transcription before migrating into the brain, and as such, directly cause the pathogenic effect as described above.

In contrast to MT1, MT2 did not change isoform preference in splicing nor did we see an impaired effect of MT2 isoform2 in our *in vitro Hes5* transcription assay that reached statistical significance. However, in this assay, the MT2 group was also not significantly different from the MT1 group(P=0.3519). Patients in Family-B and Family-C carry MT2, but both had additional variants in other genes. Patients in Family-B possessed a rare heterozygous variant in *CCDC141*, which encodes for a protein involved in embryonic GnRH neuron migration(Hutchins et al., 2016) and is a known IHH-causative gene(Turan et al., 2017). Thus, the co-occurrence of the rare variant in *CCDC141* may explain IHH in this kindred. The proband in Family-C had rare heterozygous variants in POU6F1. Although little is known about the significance of the site of the POU6F1 variants, it is possible that the combination of these variants in the closest paralogs to POU6F2 had an integrated effect to cause the IHH phenotype in this patient.

In summary, we provide evidence implicating variants in POU6F2 in the etiology of IHH with mutations in POU6F2 isoform1 directly impacting GnRH expression.

## MATERIALS AND METHODS

Human experimental protocols were approved by either the Ethics Committee of the Cukurova University Faculty of Medicine and the institutional review board of the University of Mississippi Medical Center or by the Human Research Committee at the MGH, Boston, MA. All individuals and/or their legal guardians provided written informed consent. For experiments involving mice: All procedures were performed in accordance with National Institute of Neurological Disorders and Stroke (NIH/NINDS) IACUC animal ethics guidelines (ASP-1221-20).

### Patients

Two large cohorts of IHH patients were screened for POU6F2 variants. The Cukurova cohort consisted of 416 IHH patients (nIHH, n=331 and KS, n=85) from 357 independent families recruited in Turkey. The Harvard Reproductive Endocrine Sciences Center’s IHH cohort included 677 nIHH and 632 KS patients recruited nationally and internationally. Reproductive phenotypes suggestive of IHH was deemed present if they exhibited at least one of the following IHH-related phenotypes: micropenis or cryptorchidism (boys), absent puberty by age 13 in girls and by age 14 in boys, primary amenorrhea (girls), and/or a biochemical observation of hypogonadotropic hypogonadism. The KS patients additionally had anosmia/hyposmia as determined by self-reporting and/or physical examination by administering culturally appropriate formal or informal smell tests.

### DNA sequencing and rare variant analyses

DNA samples for exome sequencing (ES) were prepared as an Illumina sequencing library, and in the second step, the sequencing libraries were enriched for the desired target using the Illumina Exome Enrichment protocol. The captured libraries were sequenced using Illumina HiSeq2000 Sequencer. The reads were mapped against UCSC (https://genome.ucsc.edu/cgi-bin/hgGateway) hg19. The variants in ES data were filtered against population polymorphism databases TR Variome(Kars et al., 2021) and gnomAD in the Cukurova cohort and against gnomAD in the Harvard cohort to obtain rare sequence variants (RSV), defined as variants with <0.001 minor allele frequency (MAF).The resulting RSVs were then screened for variants in *POU6F2* (NM_007252). The presence and segregation of significant variants within pedigrees were verified by Sanger sequencing on an Applied Biosystems PRISM 3130 auto sequencer. All animal procedures were approved by NINDS Animal Care and Use Committee and performed in accordance with NIH guidelines.

### Expression of *Pou6f2* isoforms

Total RNA was extracted from mouse adult brain, pituitary, testis, and ovaries, using Trizol reagent (Invitrogen, 15596-026) according to manufacturer’s instructions. Total RNA (1ug) was used for cDNA synthesis with oligo(dT)_16_ primer and SuperScript III Reverse Transcriptase (Invitrogen, 18080-044) following the manufacturer’s protocol. cDNAs from primary GnRH cells, generated as previously described(Kramer et al., 2000) were also analyzed. PCR analysis was performed using specific primers on *Pou6f2* exon8 (forward,5’-ACACAGACTCAGGTGGGACAA-3’) and exon9 (reverse,5’-TTCCCGGTCGTAGTTTAG-CTT-3’) or isoform2 specific primers (forward,5’-GCCATCTGCAGGTTTGAAA-3’; reverse,5’-CGTGTTGCTTTAAGCGTTTG-3’) and products compared on 2% agarose gels.

### *In vitro* splicing assay

*In silico* splicing predictions for POU domain variants were performed using online applications (Human Splicing Finder (v3.1), NetGene2, RESCUE-ESE, and Splice Site Finder by Neural Network (SSFNN)). For *in vitro* splicing assay, a mini-gene system vector was constructed using the pSPL3 plasmid (provided by Dr. C.A.Stratakis, NICHD/NIH, USA). Exon11 and flanking intronic sequences (~300bp) of the *POU6F2* gene were amplified from human gDNA and inserted into the pSPL3. Mutant constructs were generated by site-directed mutagenesis(Reikofski & Tao, 1992). After sequencing confirmation, the plasmids were transfected into HEK293FT cells (Lipofectamine^®^ LTX with PLUS^™^ reagent, Invitrogen). Cells were lysed 48hrs after transfection and total RNA extracted (Trizol Reagent). cDNA was synthesized from 1ug of RNA using oligo(dT)_16_ primer and SuperScript III Reverse Transcriptase. PCR to amplify the splicing region of exon11 of the mini-gene constructs was performed using a forward primer (SD6, on exonA of pSPL3) and an exon11 specific reverse primer. The PCR fragments from mock, WT and two mutant vectors were compared on 2% agarose gel. The sequence of PCR products was confirmed with direct Sanger sequencing after TA cloning to isolate each fragment.

### Molecular modeling

POU6F2 isoform1 and 2 were generated using C-I-TASSER(Zhang, Mortuza, He, Wang, & Zhang, 2018) from their amino acid sequences (UniProtKB codes: P78424-1 and P78424-2). Three-dimensional DNA structures were produced using w3DNA(Zheng, Lu, & Olson, 2009). POU6F2-DNA docking was simulated by HDOCK using template-free docking settings for the OCT1 DNA binding site(Yan, Tao, He, & Huang, 2020; Yan, Zhang, Zhou, Li, & Huang, 2017). As predicted POU6F2 isoform1 did not bind to this site. Since isoform1 was previously reported to bind to *Fshβ* (Yoshida et al., 2014), we tested isoform1 against the *Fshβ* protected site (5’-ATAAGCTTAAT-3’) and separately against an aligned site in the proximal promoter region of Gnrh1 (5’-AAAAGCATAGT-3’). Mutant proteins and folding free energy values for both isoforms were calculated by DynaMut(Rodrigues, Pires, & Ascher, 2018). Natural protein flexibility was detected using CABS-flex dynamics(Kuriata et al., 2018). WT vs mutant protein-DNA binding free energy values were predicted by SAMPDI(Peng, Sun, Jia, Li, & Alexov, 2018). All models were rendered using PyMOL molecular graphics software.

### *In vitro* assay for isoform2 variants

The *Hes5* gene was previously reported to be upregulated by isoform2(Fiorino et al., 2016). Thus, *in vitro* transcription assays for human POU6F2 was performed using a reporter plasmid that encodes DsRed under a Hes5 promoter (Addgene, Cat# 26868). For POU6F2 expression vectors, cDNA was introduced into pcDNA3.1(+)-IRES-GFP plasmid. Isoform1 was synthesized (GeneArt) and isoform2 and two mutant plasmids were generated by site-directed mutagenesis PCR, deleting 108bp of exon11 and subsequently introducing the mutation(Ho, Hunt, Horton, Pullen, & Pease, 1989; Reikofski & Tao, 1992). Mock or POU6F2 plasmids were co-transfected with the Hes5p-DsRed reporter plasmid into HEK293FT cells. Total protein was harvested after 3 days of culture and analyzed by Western blot (mouse anti-DsRed, Clontech; rabbit anti-POU6F2, Invitrogen, PA5-35115). Band intensities were measured using ImageJ software to quantify expression.

### *In vitro* assay for isoform1 variants

Since only isoform1 was found in GnRH cells, an *in vitro* assay for changes in GnRH expression was performed using a human GnRH cell line, FNC-B4-hTERT. ‘FNC-B4’ cells were first isolated from fetal olfactory neuroepithelium(Romanelli et al., 2004). Telomerase-mediated immortalization was performed on these cells, and the human GnRH cell line (FNC-B4-hTERT) was established(Hu et al., 2009). Cells were grown in monolayer (37°C, 5%CO2) in F-12 Coon’s modification medium (Sigma, F6636) supplemented with Penicillin-Streptomycin (Gibco, 15140-122) and 10%fetal bovine serum (Sigma, F7524). FNC-B4-hTERT cells were seeded into 6-well plates and cultured until ~80% confluency. Cells were then transfected with mock (pcDNA-3.1(+)_IRES-GFP), WT-POU6F2-isoform1, and two isoform1 mutant plasmids using FuGENE^®^ HD (Promega, E2311) following the manufacturer’s instructions. Thirty-two hrs after transfection, culture media were changed to serum free media for 16hrs prior to GnRH stimulation(Romanelli et al., 2004). Cells were treated with GnRH (0.2uM,[D-Trp^6^]-LH-RH, Sigma, L9761) for 3hrs and harvested for RNA preparation. Experiments were performed in triplicate. Total RNA was extracted using Trizol reagent. 250ng of total RNA of each group was reverse transcribed into cDNA using 50uM oligo(dT)_20_ and SuperScript III reverse transcriptase. All cDNA was stored at -20°C until analysis of GnRH transcript levels using RT-qPCR. qPCR was performed with primers specific for human Beta-Actin (*ACTB;* forward: 5’-CACCATTGGCAATGAGCGGTTC-3’; reverse, 5’-AGGTCT-TTGCGGATGTCCACGT-3’) and GnRH (*GnRH1*; forward,5’-CAACGCTTCGAATGCACCA-3’; reverse,5’-ATGTGCAACTTGGTGTAAGGATT-3’). The primer efficiency of Beta-Actin was 92.3% with an R^2^=0.9997. The primer efficiency of GnRH was 126.71% with an R^2^=0.9877, falling within a ‘good’ efficiency and amplification factor for qPCR(Taylor, Wakem, Dijkman, Alsarraj, & Nguyen, 2010). qPCR was performed using SsoAdvanced Universal SYBR^®^ Green Supermix (BioRad, 1725271) and StepOne Real-Time PCR System (Applied Biosystems). Samples amplified with the Beta-Actin primers were diluted 1:100. Samples amplified with GnRH primers were diluted 2:3. Each sample was run in triplicate.

Each group was run together on the qPCR machine which resulted in three unique runs. The average of all the automatic thresholds was taken and used to set a manual threshold. ΔΔCт was calculated to compare GnRH expression across treatment groups. This was done by first calculating the mean of the technical triplicates for each sample for each primer. The ΔCт was then calculated by taking the mean Cт value for GnRH and subtracting the mean Cт value for Beta-Actin for each sample. The ΔΔCт was calculated by subtracting the reference treatment condition (Mock) ΔCт from each of the ΔCт of the treatment conditions (WT, MT1 and MT2). Lastly, the relative expression of GnRH in each group was determined by taking 2 to the power of the negative ΔΔCт.

### Statistical analysis

Data are expressed as mean± SEM, and statistical evaluation was performed using unpaired t tests (Prism for macOS, v9.3.1). Data from *in vitro* splicing assay and transcriptional assay was normalized and represented as relative compared to control or WT group. P<0.05 was considered statistically significant. For qPCR, the Mock ΔΔCт was set to 1 and the remaining treatment conditions were adjusted accordingly to compare across experimental runs. Statistical significance between group were compared using unpaired t tests across biological triplicates.

## ACKNOWLEDGMENTS

We thank Dr. C.A. Stratakis (NICHD/NIH, USA) for providing the pSPL3 plasmid and Dr. Soo-Hyun Kim (St George’s, University of London, United Kingdom) for providing FNC-B4-hTERT cells, Lacey Plummer for the data retrieval. This study was supported by University of Mississippi Medical Center, (grant DN00305, AKT), Cukurova University (scientific research project number 11364, AKT), National Institutes of Health, National Institute of Neurological Disorders and Stroke (ZIANS-002824-30, SW), Eunice K. Shriver National Institute for Child Health and Human Development (P50HD104224-01, RB and SBS; R37 HD043341-19, R01 FD005712-04, SBS).

## COMPETING INTERESTS

The authors declare no competing interests.

## SUPPLEMENTARY FIGURES AND TABLES

**Supplementary Figure 1.**
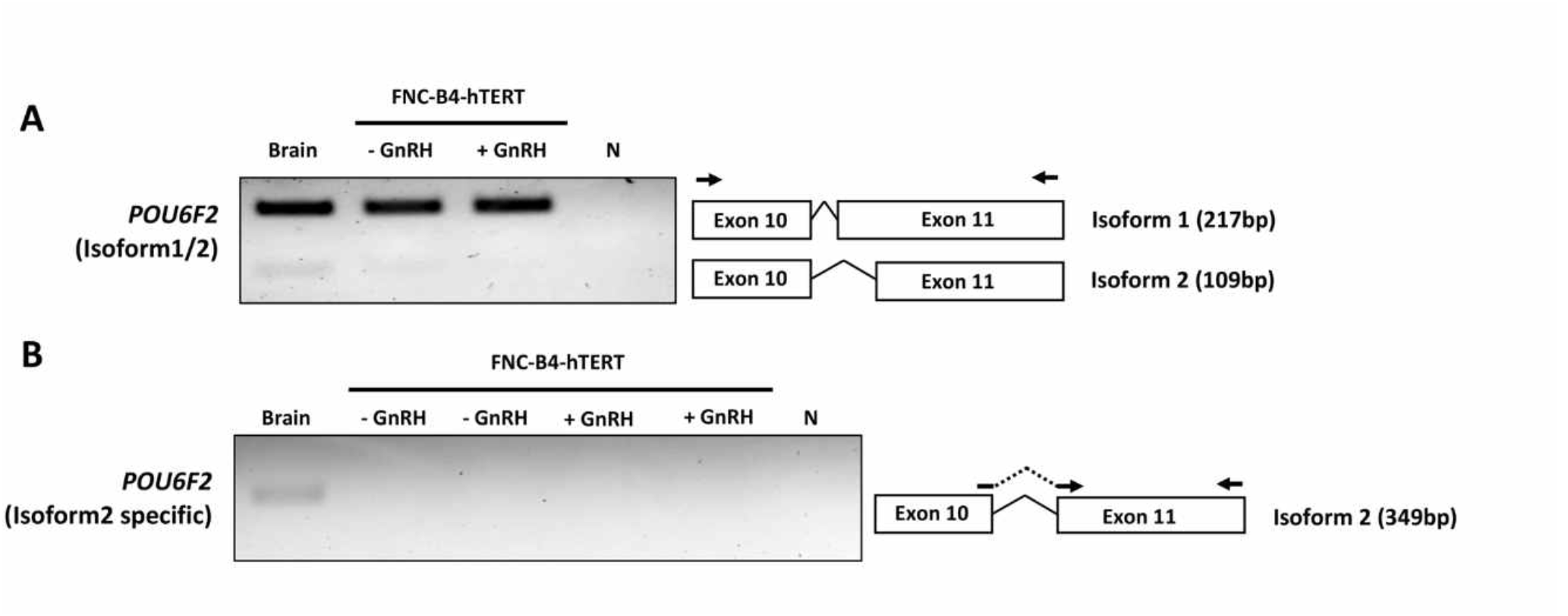
Expression of *POU6F2* isoforms in human brain and FNC-B4-hTERT cells. **(A)** RT-PCR analysis performed in human brain and immortalized GnRH cells (with or without GnRH stimulation). Top band (217 bp) shows isoform1 and bottom band (109bp) shows isoform2 which is skipping 108bp by alternative splicing on exon 11. Primers used for PCR are shown as arrows on exon 10 and 11. **(B)** RT-PCR analysis performed using isoform2 specific primers (shown as arrows on the junction of exon 10-11 and exon 11). Isoform2 (349bp) was detected in human brain but not in FNC-B4-hTERT cells.

**Supplemental Figure 2.**
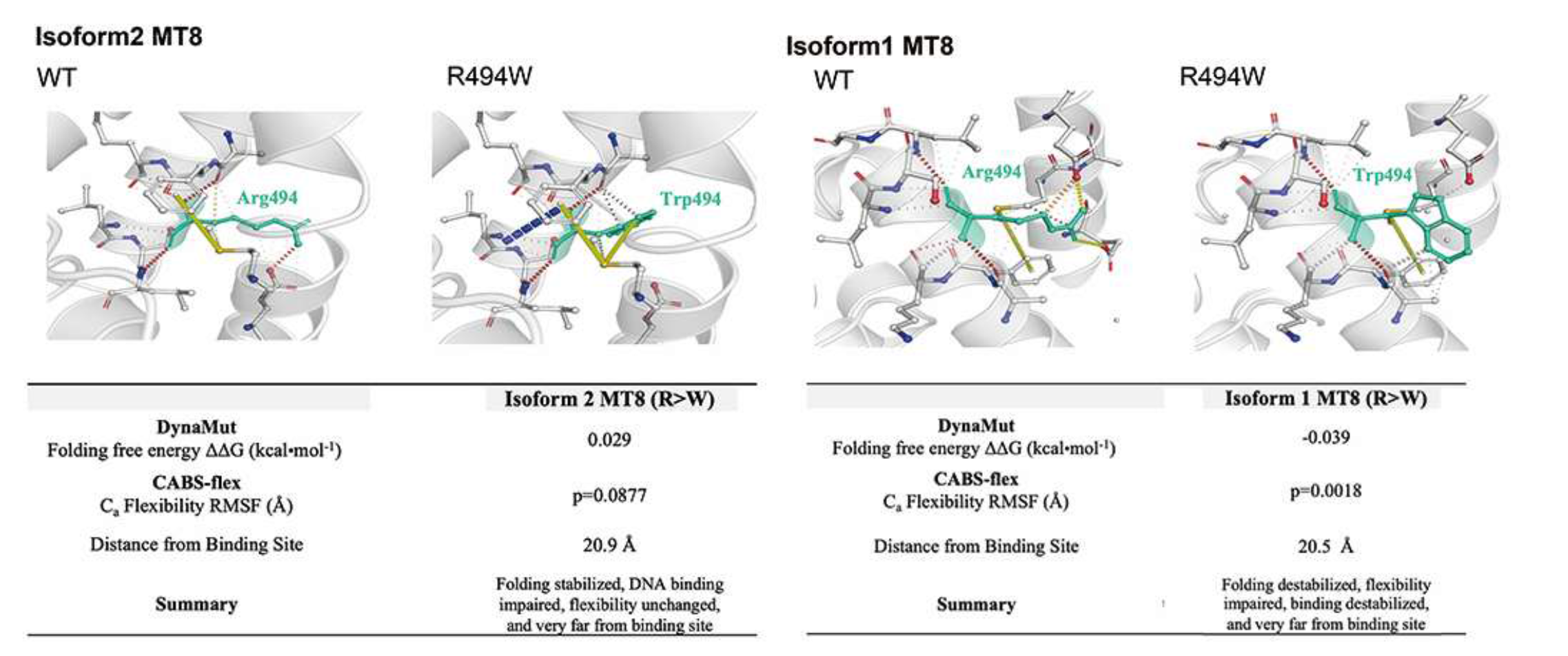
Structural analysis of IHH mutation MT8 into POU6F2 isoform1 and 2. Structural evaluation scores indicating how MT8 affects POU6F2 isoform1 and isoform2 protein folding (DynaMut) and natural protein flexibility (CABS-flex) in the individual protein structures. Characterization of stabilizing or destabilizing effects are indicated. CABS-flex values analyzed using a paired-t test.

**Supplemental Table 1.**
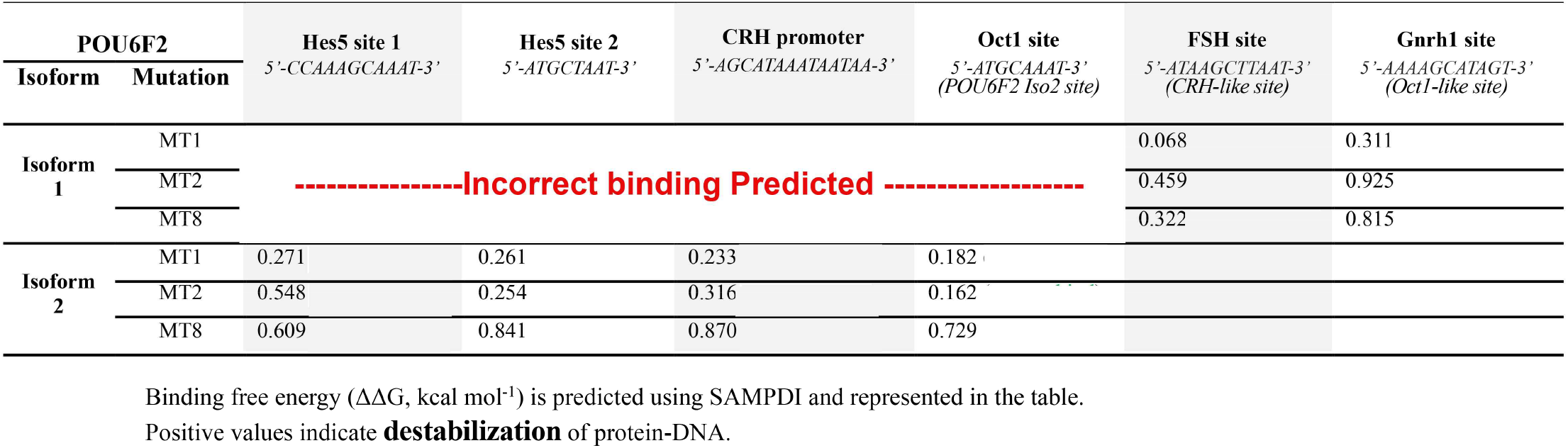
Computational analysis of variants bound to DNA targets.

## Notes

### Competing Interest Statement

The authors have declared no competing interest.

## REFERENCES

Adzhubei, I. A., Schmidt, S., Peshkin, L., Ramensky, V. E., Gerasimova, A., Bork, P., … Sunyaev, S. R. (2010). A method and server for predicting damaging missense mutations. Nat Methods, 7(4), 248–249. doi:10.1038/nmeth0410-248

Andersen, B., & Rosenfeld, M. G. (2001). POU domain factors in the neuroendocrine system: lessons from developmental biology provide insights into human disease. Endocr Rev, 22(1), 2–35. doi:10.1210/edrv.22.1.0421

Bouilly, J., Messina, A., Papadakis, G., Cassatella, D., Xu, C., Acierno, J. S., … Pitteloud, N. (2018). DCC/NTN1 complex mutations in patients with congenital hypogonadotropic hypogonadism impair GnRH neuron development. Hum Mol Genet, 27(2), 359–372. doi:10.1093/hmg/ddx408

Campbell, J. N., Macosko, E. Z., Fenselau, H., Pers, T. H., Lyubetskaya, A., Tenen, D., … Tsai, L. T. (2017). A molecular census of arcuate hypothalamus and median eminence cell types. Nat Neurosci, 20(3), 484–496. doi:10.1038/nn.4495

Clarkson, J., Han, S. Y., Piet, R., McLennan, T., Kane, G. M., Ng, J., … Herbison, A. E. (2017). Definition of the hypothalamic GnRH pulse generator in mice. Proc Natl Acad Sci U S A, 114(47), E10216–E10223. doi:10.1073/pnas.1713897114

Duittoz, A. H., Forni, P. E., Giacobini, P., Golan, M., Mollard, P., Negrón, A. L., … Wray, S. (2021). Development of the gonadotropin-releasing hormone system. J Neuroendocrinol, e13087. doi:10.1111/jne.13087

Fiorino, A., Manenti, G., Gamba, B., Bucci, G., De Cecco, L., Sardella, M., … Perotti, D. (2016). Retina-derived POU domain factor 1 coordinates expression of genes relevant to renal and neuronal development. Int J Biochem Cell Biol, 78, 162–172. doi:10.1016/j.biocel.2016.07.013

Givens, M. L., Rave-Harel, N., Goonewardena, V. D., Kurotani, R., Berdy, S. E., Swan, C. H,… Mellon, P. L. (2005). Developmental regulation of gonadotropin-releasing hormone gene expression by the MSX and DLX homeodomain protein families. J Biol Chem, 280(19), 19156–19165. doi:10.1074/jbc.M502004200

Gutierrez-Arcelus, M., Ongen, H., Lappalainen, T., Montgomery, S. B., Buil, A., Yurovsky A… Dermitzakis, E. T. (2015). Tissue-specific effects of genetic and epigenetic variation on gene regulation and splicing. PLoS Genet, 11(1), e1004958. doi:10.1371/journal.pgen.1004958

Haddad, Y., Adam, V., & Heger, Z. (2020). Ten quick tips for homology modeling of high-resolution protein 3D structures. PLoS Comput Biol, 16(4), e1007449. doi:10.1371/journal.pcbi.1007449

Herbison, A. E., Porteous, R., Pape, J. R., Mora, J. M., & Hurst, P. R. (2008). Gonadotropin-releasing hormone neuron requirements for puberty, ovulation, and fertility. Endocrinology, 149(2), 597–604. doi:10.1210/en.2007-1139

Herr, W., & Cleary, M. A. (1995). The POU domain: versatility in transcriptional regulation by a flexible two-in-one DNA-binding domain. Genes Dev, 9(14), 1679–1693. doi:10.1101/gad.9.14.1679

Ho, S. N., Hunt, H. D., Horton, R. M., Pullen, J. K., & Pease, L. R. (1989). Site-directed mutagenesis by overlap extension using the polymerase chain reaction. Gene, 77(1), 51–59. doi:10.1016/0378-1119(89)90358-2

Howard, S. R., & Dunkel, L. (2019). Delayed Puberty-Phenotypic Diversity, Molecular Genetic Mechanisms, and Recent Discoveries. Endocr Rev, 40(5), 1285–1317. doi:10.1210/er.2018-00248

Hu, Y., Guimond, S. E., Travers, P., Cadman, S., Hohenester, E., Turnbull, J. E.,… Bouloux, P. M. (2009). Novel mechanisms of fibroblast growth factor receptor 1 regulation by extracellular matrix protein anosmin-1. J Biol Chem, 284(43), 29905–29920. doi:10.1074/jbc.M109.049155

Huisman, C., Cho, H., Brock, O., Lim, S. J., Youn, S. M., Park, Y.,… Lee, J. W. (2019). Single cell transcriptome analysis of developing arcuate nucleus neurons uncovers their key developmental regulators. Nat Commun, 10(1), 3696. doi:10.1038/s41467-019-11667-y

Hutchins, B. I., Kotan, L. D., Taylor-Burds, C., Ozkan, Y., Cheng, P. J., Gurbuz, F.,… Wray, S. (2016). CCDC141 Mutation Identified in Anosmic Hypogonadotropic Hypogonadism (Kallmann Syndrome) Alters GnRH Neuronal Migration. Endocrinology, 157(5), 1956–1966. doi:10.1210/en.2015-1846

Jacobson, E. M., Li, P., Leon-del-Rio, A., Rosenfeld, M. G., & Aggarwal, A. K. (1997). Structure of Pit-1 POU domain bound to DNA as a dimer: unexpected arrangement and flexibility. Genes Dev, 11(2), 198–212. doi:10.1101/gad.11.2.198

Kars, M. E., Basak, A. N., Onat, O. E., Bilguvar, K., Choi, J., Itan, Y.,… Ozcelik, T. (2021). The genetic structure of the Turkish population reveals high levels of variation and admixture. Proc Natl Acad Sci U S A, 118(36). doi:10.1073/pnas.2026076118

Kim, K. P., Han, D. W., Kim, J., & Schöler, H. R. (2021). Biological importance of OCT transcription factors in reprogramming and development. Exp Mol Med, 53(6), 1018–1028. doi:10.1038/s12276-021-00637-4

King, J. C., & Rubin, B. S. (1995). Dynamic alterations in luteinizing hormone-releasing hormone (LHRH) neuronal cell bodies and terminals of adult rats. Cell Mol Neurobiol, 15(1), 89–106. doi:10.1007/BF02069560

Kioussi, C., Carriere, C., & Rosenfeld, M. G. (1999). A model for the development of the hypothalamic-pituitary axis: transcribing the hypophysis. Mech Dev, 81(1-2), 23–35. doi:10.1016/s0925-4773(98)00229-9

Kramer, P. R., Krishnamurthy, R., Mitchell, P. J., & Wray, S. (2000). Transcription factor activator protein-2 is required for continued luteinizing hormone-releasing hormone expression in the forebrain of developing mice. Endocrinology, 141(5), 1823–1838. doi:10.1210/endo.141.5.7452

Kramer, P. R., & Wray, S. (2000). Novel gene expressed in nasal region influences outgrowth of olfactory axons and migration of luteinizing hormone-releasing hormone (LHRH) neurons. Genes Dev, 14(14), 1824–1834.

Kukurba, K. R., Zhang, R., Li, X., Smith, K. S., Knowles, D. A., How Tan, M.,… Montgomery, S. B. (2014). Allelic expression of deleterious protein-coding variants across human tissues. PLoS Genet, 10(5), e1004304. doi:10.1371/journal.pgen.1004304

Kumar, P., Henikoff, S., & Ng, P. C. (2009). Predicting the effects of coding non-synonymous variants on protein function using the SIFT algorithm. Nat Protoc, 4(7), 1073–1081. doi:10.1038/nprot.2009.86

Kuriata, A., Gierut, A. M., Oleniecki, T., Ciemny, M. P., Kolinski, A., Kurcinski, M., & Kmiecik, S. (2018). CABS-flex 2.0: a web server for fast simulations of flexibility of protein structures. Nucleic Acids Res, 46(W1), W338–W343. doi:10.1093/nar/gky356

Leclerc, G. M., & Boockfor, F. R. (2005). Identification of a novel OCT1 binding site that is necessary for the elaboration of pulses of rat GnRH promoter activity. Mol Cell Endocrinol, 245(1-2), 86–92. doi:10.1016/j.mce.2005.10.026

Li, X., Kim, Y., Tsang, E. K., Davis, J. R., Damani, F. N., Chiang, C.,… Montgomery, S. B. (2017). The impact of rare variation on gene expression across tissues. Nature, 550(7675), 239–243. doi:10.1038/nature24267

Louden, E. D., Poch, A., Kim, H. G., Ben-Mahmoud, A., Kim, S. H., & Layman, L. C. (2021). Genetics of hypogonadotropic Hypogonadism-Human and mouse genes, inheritance, oligogenicity, and genetic counseling. Mol Cell Endocrinol, 534, 111334. doi:10.1016/j.mce.2021.111334

Miao, Y., Li, C., Guo, J., Wang, H., Gong, L., Xie, W., & Zhang, Y. (2019). Identification of a novel somatic mutation of POU6F2 by whole-genome sequencing in prolactinoma. Mol Genet Genomic Med, 7(12), e1022. doi:10.1002/mgg3.1022

Peng, Y., Sun, L., Jia, Z., Li, L., & Alexov, E. (2018). Predicting protein-DNA binding free energy change upon missense mutations using modified MM/PBSA approach: SAMPDI webserver. Bioinformatics, 34(5), 779–786. doi:10.1093/bioinformatics/btx698

Pereira, J. H., & Kim, S. H. (2009). Structure of human Brn-5 transcription factor in complex with CRH gene promoter. J Struct Biol, 167(2), 159–165. doi:10.1016/j.jsb.2009.05.003

Pitteloud, N., Hayes, F. J., Dwyer, A., Boepple, P. A., Lee, H., & Crowley, W. F., Jr. (2002). Predictors of outcome of long-term GnRH therapy in men with idiopathic hypogonadotropic hypogonadism. J Clin Endocrinol Metab, 87(9), 4128–4136. doi:10.1210/jc.2002-020518

Reikofski, J., & Tao, B. Y. (1992). Polymerase chain reaction (PCR) techniques for site-directed mutagenesis. Biotechnol Adv, 10(4), 535–547. doi:10.1016/0734-9750(92)91451-j

Reményi, A., Tomilin, A., Pohl, E., Lins, K., Philippsen, A., Reinbold, R.,… Wilmanns, M. (2001). Differential dimer activities of the transcription factor Oct-1 by DNA-induced interface swapping. Mol Cell, 8(3), 569–580. doi:10.1016/s1097-2765(01)00336-7

Renault, C. H., Aksglaede, L., Wojdemann, D., Hansen, A. B., Jensen, R. B., & Juul, A. (2020). Minipuberty of human infancy - A window of opportunity to evaluate hypogonadism and differences of sex development? Ann Pediatr Endocrinol Metab, 25(2), 84–91. doi:10.6065/apem.2040094.047

Richards, S., Aziz, N., Bale, S., Bick, D., Das, S., Gastier-Foster, J.,… Committee, A. L. Q. A. (2015). Standards and guidelines for the interpretation of sequence variants: a joint consensus recommendation of the American College of Medical Genetics and Genomics and the Association for Molecular Pathology. Genet Med, 17(5), 405–424. doi:10.1038/gim.2015.30

Rodrigues, C. H., Pires, D. E., & Ascher, D. B. (2018). DynaMut: predicting the impact of mutations on protein conformation, flexibility and stability. Nucleic Acids Res, 46(W1), W350–W355. doi:10.1093/nar/gky300

Romanelli, R. G., Barni, T., Maggi, M., Luconi, M., Failli, P., Pezzatini, A.,… Vannelli, G. B. (2004). Expression and function of gonadotropin-releasing hormone (GnRH) receptor in human olfactory GnRH-secreting neurons: an autocrine GnRH loop underlies neuronal migration. J Biol Chem, 279(1), 117–126. doi:10.1074/jbc.M307955200

Ryan, A. K., & Rosenfeld, M. G. (1997). POU domain family values: flexibility, partnerships, and developmental codes. Genes Dev, 11(10), 1207–1225. doi:10.1101/gad.11.10.1207

Saengkaew, T., Ruiz-Babot, G., David, A., Mancini, A., Mariniello, K., Cabrera, C. P.,… Howard, S. R. (2021). Whole exome sequencing identifies deleterious rare variants in CCDC141 in familial self-limited delayed puberty. NPJ Genom Med, 6(1), 107. doi:10.1038/s41525-021-00274-w

Sanz, E., Quintana, A., Deem, J. D., Steiner, R. A., Palmiter, R. D., & McKnight, G. S. (2015). Fertility-regulating Kiss1 neurons arise from hypothalamic POMC-expressing progenitors. J Neurosci, 35(14), 5549–5556. doi:10.1523/JNEUROSCI.3614-14.2015

Simonian, S. X., & Herbison, A. E. (2001). Regulation of gonadotropin-releasing hormone (GnRH) gene expression during GnRH neuron migration in the mouse. Neuroendocrinology, 73(3), 149–156. doi:10.1159/000054631

Strande, N. T., Riggs, E. R., Buchanan, A. H., Ceyhan-Birsoy, O., DiStefano, M., Dwight, S. S.,… Berg, J. S. (2017). Evaluating the Clinical Validity of Gene-Disease Associations: An Evidence-Based Framework Developed by the Clinical Genome Resource. Am J Hum Genet, 100(6), 895–906. doi:10.1016/j.ajhg.2017.04.015

Taylor, S., Wakem, M., Dijkman, G., Alsarraj, M., & Nguyen, M. (2010). A practical approach to RT-qPCR-Publishing data that conform to the MIQE guidelines. Methods, 50(4), S1–5. doi:10.1016/j.ymeth.2010.01.005

Teilum, K., Olsen, J. G., & Kragelund, B. B. (2011). Protein stability, flexibility and function. Biochim Biophys Acta, 1814(8), 969–976. doi:10.1016/j.bbapap.2010.11.005

Turan, I., Hutchins, B. I., Hacihamdioglu, B., Kotan, L. D., Gurbuz, F., Ulubay, A.,… Topaloglu, A. K. (2017). CCDC141 Mutations in Idiopathic Hypogonadotropic Hypogonadism. J Clin Endocrinol Metab, 102(6), 1816–1825. doi:10.1210/jc.2016-3391

Turton, J. P., Reynaud, R., Mehta, A., Torpiano, J., Saveanu, A., Woods, K. S.,… Dattani, M. T. (2005). Novel mutations within the POU1F1 gene associated with variable combined pituitary hormone deficiency. J Clin Endocrinol Metab, 90(8), 4762–4770. doi:10.1210/jc.2005-0570

Wierman, M. E., Xiong, X., Kepa, J. K., Spaulding, A. J., Jacobsen, B. M., Fang, Z.,… Ojeda, S. R. (1997). Repression of gonadotropin-releasing hormone promoter activity by the POU homeodomain transcription factor SCIP/Oct-6/Tst-1: a regulatory mechanism of phenotype expression? Mol Cell Biol, 17(3), 1652–1665. doi:10.1128/mcb.17.3.1652

Wolfe, A., Kim, H. H., Tobet, S., Stafford, D. E., & Radovick, S. (2002). Identification of a discrete promoter region of the human GnRH gene that is sufficient for directing neuron-specific expression: a role for POU homeodomain transcription factors. Mol Endocrinol, 16(3), 435–449. doi:10.1210/mend.16.3.0780

Xu, C., Messina, A., Somm, E., Miraoui, H., Kinnunen, T., Acierno, J., Jr.,… Pitteloud, N. (2017). KLB, encoding beta-Klotho, is mutated in patients with congenital hypogonadotropic hypogonadism. EMBO Mol Med, 9(10), 1379–1397. doi:10.15252/emmm.201607376

Yan, Y., Tao, H., He, J., & Huang, S. Y. (2020). The HDOCK server for integrated protein-protein docking. Nat Protoc, 15(5), 1829–1852. doi:10.1038/s41596-020-0312-x

Yan, Y., Zhang, D., Zhou, P., Li, B., & Huang, S. Y. (2017). HDOCK: a web server for protein-protein and protein-DNA/RNA docking based on a hybrid strategy. Nucleic Acids Res, 45(W1), W365–W373. doi:10.1093/nar/gkx407

Yoshida, S., Ueharu, H., Higuchi, M., Horiguchi, K., Nishimura, N., Shibuya, S.,… Kato, Y. (2014). Molecular cloning of rat and porcine retina-derived POU domain factor 1 (POU6F2) from a pituitary cDNA library. J Reprod Dev, 60(4), 288–294. doi:10.1262/jrd.2014-023

Zhang, C., Mortuza, S. M., He, B., Wang, Y., & Zhang, Y. (2018). Template-based and free modeling of I-TASSER and QUARK pipelines using predicted contact maps in CASP12. Proteins, 86 Suppl 1, 136–151. doi:10.1002/prot.25414

Zheng, G., Lu, X. J., & Olson, W. K. (2009). Web 3DNA--a web server for the analysis, reconstruction, and visualization of three-dimensional nucleic-acid structures. Nucleic Acids Res, 37(Web Server issue), W240–246. doi:10.1093/nar/gkp358

Zhou, H., Yoshioka, T., & Nathans, J. (1996). Retina-derived POU-domain factor-1: a complex POU-domain gene implicated in the development of retinal ganglion and amacrine cells. J Neurosci, 16(7), 2261–2274.

Zhu, J., Choa, R. E., Guo, M. H., Plummer, L., Buck, C., Palmert, M. R.,… Chan, Y. M. (2015). A shared genetic basis for self-limited delayed puberty and idiopathic hypogonadotropic hypogonadism. J Clin Endocrinol Metab, 100(4), E646–654. doi:10.1210/jc.2015-1080

